# THE SPATIAL REGULATION OF CONDENSIN ACTIVITY IN CHROMOSOME CONDENSATION

**DOI:** 10.1101/849273

**Authors:** Rebecca Lamothe, Lorenzo Costantino, Douglas E Koshland

## Abstract

Condensin mediates chromosome condensation, which is essential for proper chromosome segregation during mitosis. Prior to anaphase of budding yeast, the ribosomal DNA (RDN) condenses to a thin loop that is distinct from the rest of the chromosomes. We provide evidence that the establishment and maintenance of this RDN condensation require the regulation of condensin by Cdc5p (polo) kinase. We show that Cdc5p is recruited to the site of condensin binding in the RDN by cohesin, a complex related to condensin. Cdc5p and cohesin prevent condensin from misfolding the RDN into an irreversibly decondensed state. From these and other observations, we propose that the spatial regulation of Cdc5p by cohesin modulates condensin activity to ensure proper RDN folding into a thin loop. This mechanism may be evolutionarily conserved, promoting the thinly condensed constrictions that occur at centromeres and RDN of mitotic chromosomes in plants and animals.

## Introduction

Mitotic chromosomes have two structural features, sister chromatid cohesion, and chromosome condensation, which are essential for their proper inheritance during cell division. Sister chromatid cohesion is established concurrently with DNA replication and is maintained until the onset of anaphase. In contrast, condensation is established in mitotic prophase and persists through anaphase until the end of mitotic division. As a result, cohesion and condensation coexist from prophase until the onset of anaphase, a period henceforth referred to as mid-M. Perturbing proper condensation or cohesion leads to aneuploidy and chromosome rearrangements, hallmarks of cancer, birth defects and several genetic disorders (Hassler et al., 2018). Over the last two decades, scientists have made significant progress in identifying the protein complexes that mediate cohesion and condensation and understanding their cell cycle regulation. However, the potential interplay between these complexes has remained largely uninvestigated.

Cohesion and condensation are mediated respectively by cohesin and condensin, two related protein complexes in the SMC (structural maintenance of chromosomes) family (Figure 1A) (Hassler et al., 2018). Cohesin intermolecularly tethers DNA of the sister chromatids, resulting in cohesion, while condensin promotes condensation by tethering DNA intramolecularly. Condensin is also thought to promote condensation by inducing supercoils and by actively extruding DNA to form loops (Ganji et al., 2018; Hagstrom et al., 2002; Kimura et al., 2001; Kimura and Hirano, 1997; St-Pierre et al., 2009). In principle, the existence of distinct SMC complexes dedicated to cohesion and condensation would allow these processes to occur independently. Consistent with this conclusion, mammalian chromosomes can condense in the absence of cohesin (Losada et al., 1998; Sonoda et al., 2001).

**Figure 1:**
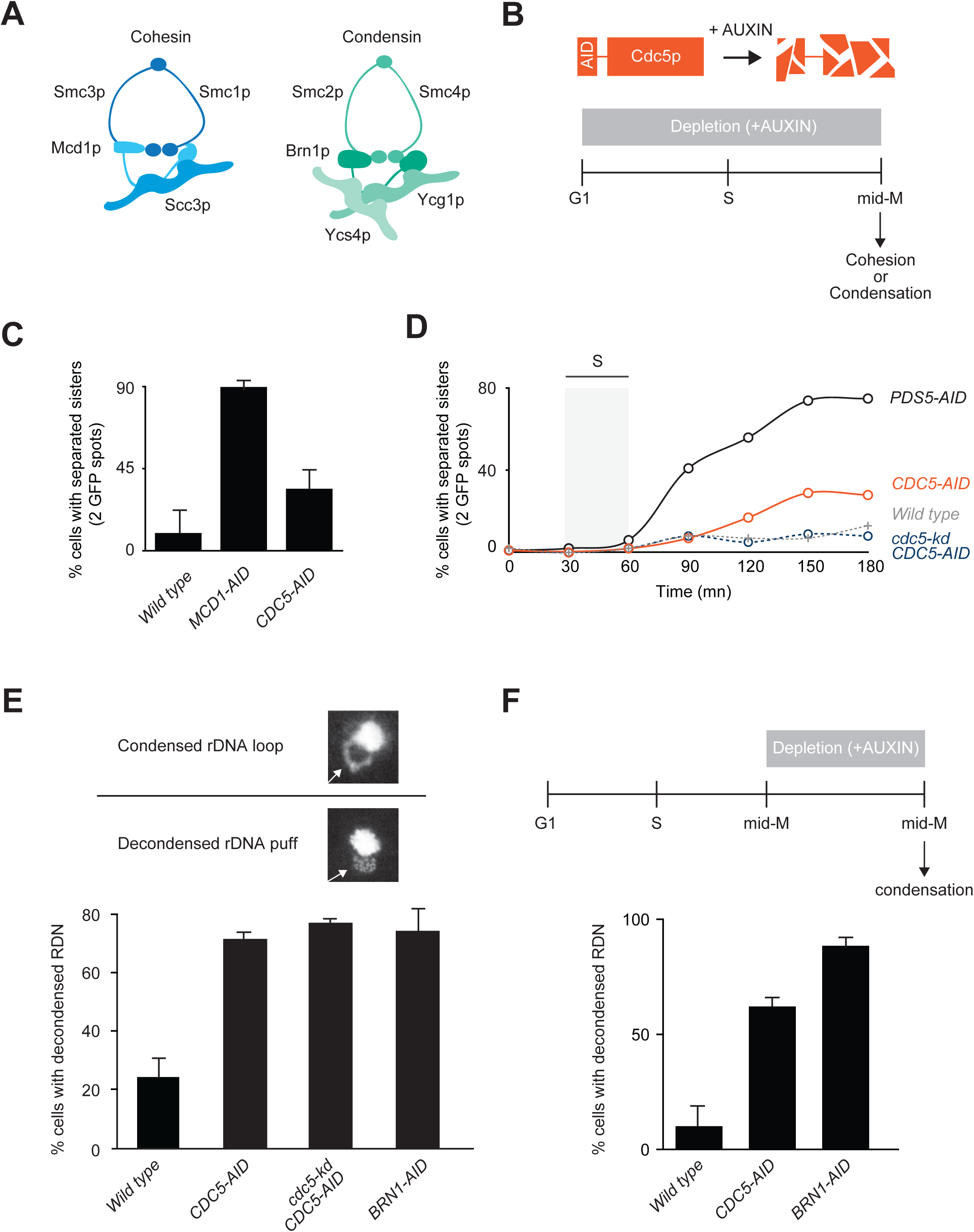
*The polo kinase Cdc5p is essential for mid-M chromosome condensation* **A.** Schematic of cohesin and condensin. Cohesin is comprised of a Smc1p/Smc3p heterodimer joined at their head domain through Mcd1p. Mcd1p is likewise bound to Scc3p resulting in the cohesin holocomplex. Condensin is comprised of a Smc2p/Smc4p heterodimer linked by Brn1p. Brn1p is bound to Ycg1p and Ycs4p. Together, these proteins form the pentameric condensin complex. **B.** *Top:* Schematic of the auxin-induced degron (AID) system to study functions of essential proteins. Proteins of interest, like Cdc5p, were fused to a 3V5-AID tag that allows for wild-type functions in the absence of auxin. Upon addition of auxin, 3V5-AID-tagged proteins were rapidly degraded in a proteasome-mediated manner. *Bottom:* ‘Staged mid-M’ regimen used to assess strains function in mid-M. Cultures were synchronized in G1 and released in media containing auxin and nocodazole, to depleted AID-tagged protein and arrest cells in mid-M. **C.** Depletion of Cdc5p results in a moderate cohesion defect. Cells were treated as in 1B and cohesion assayed in mid-M through a GFP spot assay. In these experiments, LacO arrays are integrated at LYS4, a centromere-distal arm site. Wild type, Mcd1p depleted, and Cdc5p depleted cells were scored for sister chromatid cohesion. The average percentage of separated sister chromatids from three independent experiments (100 cells) are reported, and error bars represent standard deviation. **D.** Depletion of Cdc5p results in a moderate cohesion maintenance defect. Wild type, Pds5p depleted, Cdc5p depleted, and Cdc5 kinase-dead (kd) expressing in combination with depletion of Cdc5p-AID cells were treated as in B and cohesion assayed at 15mn intervals after release from G1 to mid-M. The average percentage of separated sister chromatids from two independent experiments (100 cells) are reported. **E.** Depletion of Cdc5p results in condensation loss. *Top:* Example of rDNA locus (RDN) morphologies assayed. RDN is either assayed as a condensed loop or decondensed puff. The percentage of cells displaying decondensed RDN puffs is quantified. *Bottom:* Wild type, Cdc5p depleted, Cdc5 kinase-dead (kd) expressing in combination with depletion of Cdc5p-AID, and Brn1p depleted cells were treated as in B and processed to make chromosome spreads to score RDN condensation (see Materials and Methods). Average of three experiments scoring 200 cells are shown and error bars represent the standard deviation. **F.** Depletion of Cdc5p results in loss of maintenance of condensation. *Top:* Cells were arrested in mid-M using nocodazole. The arrest was confirmed by analysis of bud morphology (>95% large budded cells). Auxin was added to ensure Cdc5p-AID degradation. *Bottom:* Wild type, Cdc5p depleted, and Brn1p depleted cells were processed to assess condensation as described in E.

The independence of cohesion and condensation also seems to be supported by studies of their regulation. The establishment of cohesion in S phase is regulated by Eco1p-dependent acetylation of cohesin (Rolef Ben-Shahar et al., 2008; Unal et al., 2008), while its maintenance is mediated by interactions with Pds5p, a cohesin auxiliary factor (Noble et al., 2006). In contrast, the phosphorylation of condensin by different kinases mediates the establishment and maintenance of condensation. Cyclin-dependent kinases (CDKs) are required to establish condensation in prophase (Kimura et al., 1998). Polo kinase and Aurora kinase are important for the maintenance of condensation after the onset of anaphase when CDK activity diminishes (Lavoie et al., 2002, 2004; Robellet et al., 2015; St-Pierre et al., 2009). These observations suggest cohesion and condensation are mediated by distinct protein complexes with distinct regulatory circuits. However, cohesin inactivation abrogates condensation as well as cohesion in mitosis of budding yeast, meiosis of fission yeast, and mouse oocytes and sperm (Ding et al., 2006; Guacci and Koshland, 2012; Guacci et al., 1997b; Novak et al., 2008). A subsequent study in budding yeast showed that cohesin is not required for condensation per se, but rather prevents condensin from permanently misfolding chromosomes into an irreversibly decondensed state (Lavoie et al., 2002). How cohesin regulated condensin in mid-M remained a mystery.

A candidate for mediating this regulation was polo kinase. In budding yeast, polo kinase (Cdc5p) binds to cohesin associated regions (CARs) on chromosomes, and the binding is dependent upon cohesin (Pakchuen et al., 2016). Cdc5p also is required for mid-M condensation and phosphorylates condensin in mid-M (Archambault et al., 2015; Pakchuen et al., 2016; St-Pierre et al., 2009; Walters et al., 2014). Here we show that Cdc5p is required for both the establishment and maintenance of condensation of the one megabase ribosomal DNA (RDN) in mid-M, likely by promoting condensin activity to make chromosome folds. We provide additional evidence that suggests cohesin recruits Cdc5p to the RDN, facilitating the activation of adjacently bound condensin. This spatial and temporal activation of condensin promotes RDN condensation into a thin loop and prevents condensin-mediated RDN misfolding.

## Results

### The polo kinase Cdc5p is essential for mid-M chromosome condensation

The phenotypes of the different temperature-sensitive *cdc5* alleles vary significantly (St-Pierre et al., 2009; Walters et al., 2014). With this in mind, we generated a more stringent loss-of-function *cdc5* allele to study the role of Cdc5p in cohesion and condensation. To assess how essential is Cdc5p function in these processes, we compared Cdc5p depletion with depletion of cohesin and condensin. We took advantage of the auxin-induced degron (AID) system to generate a *cdc5* allele (Cdc5p-AID) (Nishimura et al., 2009). Briefly, in this system, the fusion of Cdc5p with an AID tag allows for near wild-type function in the absence of auxin. However, in the presence of auxin, tagged proteins are targeted by the E3 ligase TIR1 for rapid ubiquitin-dependent degradation (Figure 1B, top). We used the AID system to also deplete cohesin (Mcd1p-AID) and condensin (Brn1-AID). After depletion, the AID proteins were undetectable by western blot and all strains were inviable (Figure 1—figure supplement 1). Indicated yeast strains were first synchronized in G1, then released into media with auxin to deplete the AID-tagged protein and nocodazole to arrest cells in mid-M (Figure 1B, bottom). Using this protocol (referred to as staged mid-M) cells were depleted of the AID-tagged protein from G1 to mid-M. We proceeded in assessing sister chromatid cohesion and condensation.

We measured sister chromatid cohesion using a well-characterized GFP dot assay (Straight et al., 1996). In this system, a lacO array is integrated at a chromosomal locus and bound by lacI-GFP. If proper sister chromatid cohesion is present, the two sister chromatids are held together, and the two arrays appear as one GFP spot. Conversely, if cohesion is absent, two distinct GFP spots are observed (Guacci et al., 1997a; Michaelis et al., 1997). We tested wild type, Cdc5p-AID depleted and Mcd1p-AID depleted cells. Mcd1p depletion served as positive control for the abrogation of cohesion. Cells lacking Cdc5p-AID had a three-fold decrease in sister chromatid cohesion compared to wild-type cells (Figure 1C). This defect was significantly less than the nine-fold decrease in cohesion in cells depleted for Mcd1p-AID. Thus, Cdc5p is important, but not essential for sister chromatid cohesion.

We performed a time-course experiment for cohesion to assess whether Cdc5p functions in the establishment or maintenance of cohesion. Depletion of factors that are important for the establishment of cohesion resulted in a high level of separated sister chromatids from the end of S-phase till mid-M. While depletion of factors that are involved in the maintenance of cohesion resulted in a gradual loss of cohesion that starts at the end of S-phase and peaks at mid-M (Eng et al., 2014; Jin et al., 2009). We assessed cohesion on wild-type, Cdc5p depleted, and Pds5p depleted cells and assayed cohesion at 15-minute intervals from G1 to mid-M. Pds5p is a factor required for the maintenance of sister chromatid cohesion after S phase. Under this regimen, Cdc5p-AID depleted cells presented a weaker defect in cohesion maintenance compared to cells depleted of Pds5p-AID (Figure 1D). We concluded that Cdc5p is not involved in establishing cohesion but is partially required for cohesion maintenance. Furthermore, cells that expressed a kinase-dead version of Cdc5p in combination with depletion of the endogenous copy of Cdc5p did not present the maintenance defect and were indistinguishable from wild type (Figure 1D). Therefore, the partial requirement of Cdc5p in cohesion maintenance is independent of its kinase activity.

Using the same regimen described above, we also interrogated the role of Cdc5p in mid-M chromosome condensation by assessing the morphology of the ∼1.4 megabases ribosomal DNA model locus (RDN). The RDN contains 75-100 tandem repeats of the 9.1 kb rDNA and undergoes a series of distinct, cell cycle-dependent morphological changes (Figure 1E, top) (Guacci et al., 1994). In interphase, the RDN is a puff that is segregated to the periphery of the bulk chromosomal mass. In contrast, in mid-M, the RDN forms a condensed loop. This dramatic cytological RDN change is a tractable assay for condensation. In contrast, the rest of the genome only undergoes two-fold compaction that is only measurable by laborious FISH or GFP assays (Guacci et al., 1994; Vas et al., 2007). In mid-M arrested wild type, around 25% of cells fail to condense the RDN. In contrast, approximately 75% of Cdc5p-AID depleted cells failed to condense the RDN, similarly to depletion of a core condensin subunit, Brn1p-AID (Figure 1E, bottom; Figure 1—figure supplement 2A). Therefore, Cdc5p-AID depletion caused a severe condensation defect. Ectopic expression of a kinase-dead Ccd5p did not rescue the condensation defect of cells depleted for Cdc5p-AID. Thus, in contrast to cohesion, Cdc5p and its kinase activity play a crucial role in RDN condensation in mid-M.

To assess the role of Cdc5p in the establishment of condensation, we performed a time course following RDN condensation in Cdc5p-AID depleted cells. 90 minutes is the earliest time point after release from G1 where we could observe condensed RDN loops. Cells depleted for Cdc5p did not establish condensation at 90 minutes or later (Figure 1— figure supplement 2B). Instead, control cells depleted for the maintenance factor Pds5p showed condensed loops at 90 minutes that were lost over time. Therefore, Cdc5p is required for the establishment of condensation. To assess whether Cdc5p also functions in condensation maintenance, we modified the auxin regiment of *CDC5-AID* cells. We first arrested them in mid-M by adding nocodazole, allowing RDN condensation to occur (Figure 1—figure supplement 3) and then we added auxin to deplete Cdc5p-AID (Figure 1F, top). We observed a dramatic loss of RDN condensation in cells depleted of Cdc5p-AID in mid-M like that observed in cells depleted for Brn1p-AID (Figure 1F, bottom). Therefore, Cdc5p is an essential regulator for the establishment and maintenance of condensation in mid-M.

### Cdc5p stimulates condensin-dependent RDN condensation

How does Cdc5p regulate condensation? Insights come from further examining the RDN condensation defects. Puffed RDN in mid-M can result from either an unfolded state such as is normally seen in G1 or from a misfolded state (Lavoie et al., 2002). The unfolded state is still competent to be folded, while the misfolded state is irreversibly trapped in a decondensed manner (Figure 2A). These two states were previously distinguished using a simple genetic test referred to here as add later test (Figure 2B, top and (Lavoie et al., 2002). Briefly, temperature-sensitive condensation factors were inactivated from G1 to mid-M preventing condensation, then re-activated in mid-M and the reversibility of the condensation defect was assessed. With this test, we showed delayed activation of condensin restores RDN condensation, while delayed activation of cohesin did not (Figure 2B, bottom; Lavoie et al., 2002). Therefore, condensin inactivation results in an unfolded state, while cohesin inactivation results in an irreversibly misfolded state.

**Figure 2:**
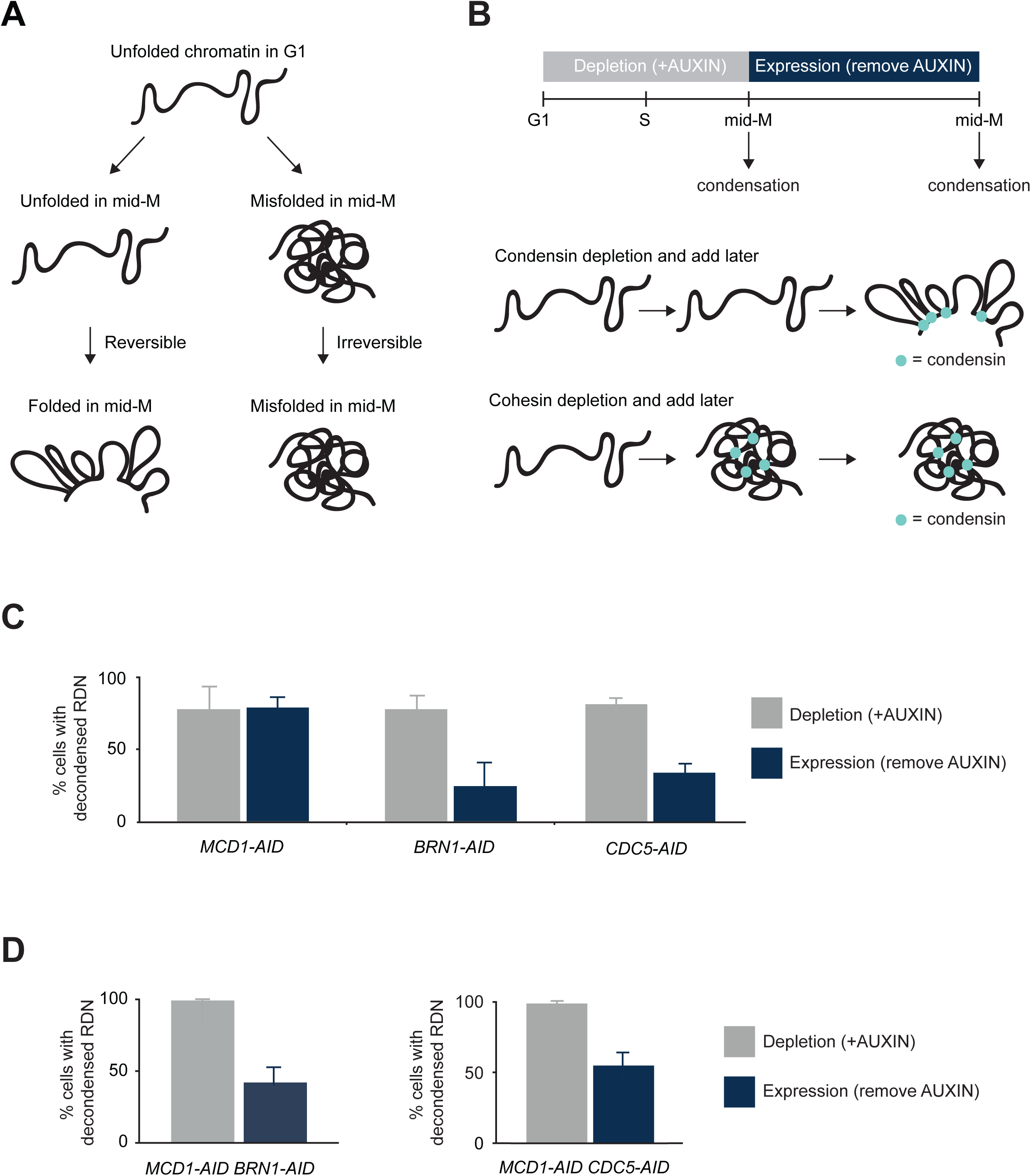
*Cdc5p stimulates condensin-dependent RDN condensation* **A.** Schematic of defects in condensation. RDN is in a puffed unfolded state in G1. Depletion of condensation factors results in a puffed RDN that can either be in an unfolded state or in a misfolded state. The unfolded state is reversible, once the factor is expressed again the RDN can be folded. Misfolded state is irreversible, even after expressing again the condensation factor the RDN remains misfolded. **B.** Schematic of ‘add later’ genetic test setup. *Top:* Indicated cultures were arrested in G1 (>95% schmooed morphology), and AID-tagged proteins (Mcd1p-AID, Brn1p-AID, Cdc5p-AID) were depleted by the addition of auxin. G1 arrest was relieved by resuspending in media with auxin and nocodazole to arrest cells in mid-M. Cells were then resuspended in media containing only nocodazole for two hours to allow the expression of the previously depleted AID-tagged proteins. *Bottom:* Depletion of condensin resulted in unfolded RDN that was folded upon later expression of condensin. Depletion of cohesin resulted in misfolded RDN that stayed misfolded upon later expression of cohesin. **C.** Cdc5p depletion results in a reversible unfolded RDN state. **‘**Add later’ genetic test results from *MCD1-AID, BRN1-AID*, and *CDC5-AID* strains were treated as described in B and scored for RDN condensation. The percentage of cells displaying puffed RDN (decondensed) is quantified. Average of two experiments scoring 300 cells are shown and error bars represent standard deviation. **D.** Condensin and Cdc5p promote misfolding of the RDN in the absence of cohesin. ‘Add-later’ genetic test results from MCD1-AID/BRN1-AID and MCD1-AID/CDC5-AID strains were treated as described in B and scored for RDN condensation. The percentage of cells displaying decondensed puffed RDN (decondensed) is quantified. Average of three experiments scoring 300 cells are shown and error bars represent standard deviation.

Cdc5p depletion can result in an unfolded state like condensin, or a misfolded state like cohesin. We performed the add later experiments using our AID system to deplete Mcd1p-AID, Brn1p-AID, and Cdc5p-AID (Figure 2—figure supplement 1). As expected, depletion of any of these three proteins between G1 and mid-M prevented the appearance of folded RDN (Figure 2C, gray bars). When Cdc5p-AID was expressed in Cdc5p-AID depleted cells, the RDN condensed (Figure 2C, blue bars). The same result was observed when Brn1p-AID was expressed in Brn1p-AID depleted cells. Conversely, when Mcd1p-AID was expressed in Mcd1p-AID depleted cells, the RDN remained decondensed. These results show that depletion of Cdc5p-AID, like depletion of condensin, results in a reversible unfolded RDN state, rather than irreversibly misfolded like depletion of cohesin.

We hypothesized that the RDN misfolding that occurred after cohesin depletion was condensin and Cdc5p dependent. If this model was correct, misfolding of the RDN due to cohesin depletion could be prevented by co-inactivation of condensin or any factor that stimulates condensin activity. To test this prediction, we depleted and added later two components together (Figure 2—figure supplement 1A). When we co-depleted and added later cohesin and condensin we restored condensation, thus preventing RDN misfolding caused by the absence of cohesin alone (Figure 2D; Lavoie et al., 2002). When we depleted and added later cohesin and Cdc5p-AID together, we also prevented RDN misfolding caused by the absence of cohesin (Figure 2D). To explain these genetic results, we propose that Cdc5p is a factor that stimulates condensin activity. When cohesin is present, Cdc5p stimulates condensin to promote proper RDN folding generating condensation. When cohesin is missing, Cdc5p inappropriately stimulates condensin resulting in a misfolded decondensed RDN.

This model predicts that Cdc5p should phosphorylate condensin with or without cohesin but the pattern should change in the absence of cohesin. To test this model we staged cells in mid-M with or without cohesin. We measured the level of phosphorylation of the condensin subunit Ycg1p, which possesses the highest number of Cdc5p phospho-sites (St-Pierre et al., 2009), using a Phos-tag gel that slows the migration of phosphorylated proteins. Phosphorylated condensin was observed both in the presence or absence of cohesin as predicted (Figure 2—figure supplement 1B). Furthermore, the phosphorylation pattern was different in the absence of cohesin: we observed a higher amount of phosphorylated condensin that migrated slower, indicating either more sites being phosphorylated and/or altered phosphorylation pattern. These patterns of Ycg1 phosphorylation are consistent with our model that cohesin is required to achieve proper phosphorylation of condensin.

### Cohesin-dependent recruitment of Cdc5p to the RDN promotes its binding to condensin in the RDN

To dissect the molecular mechanism for how Cdc5p, cohesin, and condensin regulate condensation, we examined their genomic localization by ChIP-seq. We analyzed the binding of Cdc5p, cohesin, and condensin in wild-type and Cdc5p-, cohesin-, and condensin-depleted cells. We also performed an important ChIP-seq control in yeast by immunoprecipitating GFP from GFP-expressing cells. This ChIP-seq provided us with the background signal of genomic regions prone to entrapment by ChIP-seq in the absence of a real signal (Teytelman et al., 2013).

Using the staged mid-M regimen, we first assessed the binding of condensin (Ycg1p-Myc), cohesin (Mcd1p), and Cdc5p (Cdc5p-Flag) in wild-type cells (Figure 3). We observed Ycg1p enrichment in a large pericentric region peaking at centromeres and within the end of the non-transcribed region of the RDN (NTS1), corroborating previous results (Figure 3; Leonard et al., 2015). Additional peaks of enrichment for Ycg1-Myc that were previously reported (D’Ambrosio et al., 2008) were also observed along the chromosome arms, but they were likely false positives because they coincided with peaks in GFP-control (Figure 3A). Mcd1p was bound to centromeres, pericentric regions, cohesin associated regions (CARs) along chromosome arms, and a region between the two non-transcribed spacers (NTS1 and NTS2) in the rDNA repeat (Figure 3; Glynn et al., 2004; Laloraya et al., 2000; Pakchuen et al., 2016). Finally, the binding pattern of Cdc5p-Flag mirrored cohesin binding genome-wide (Figure 3; Pakchuen et al., 2016; Rossio et al., 2010). Importantly, we showed that Cdc5p co-localizes with cohesin on the RDN.

**Figure 3:**
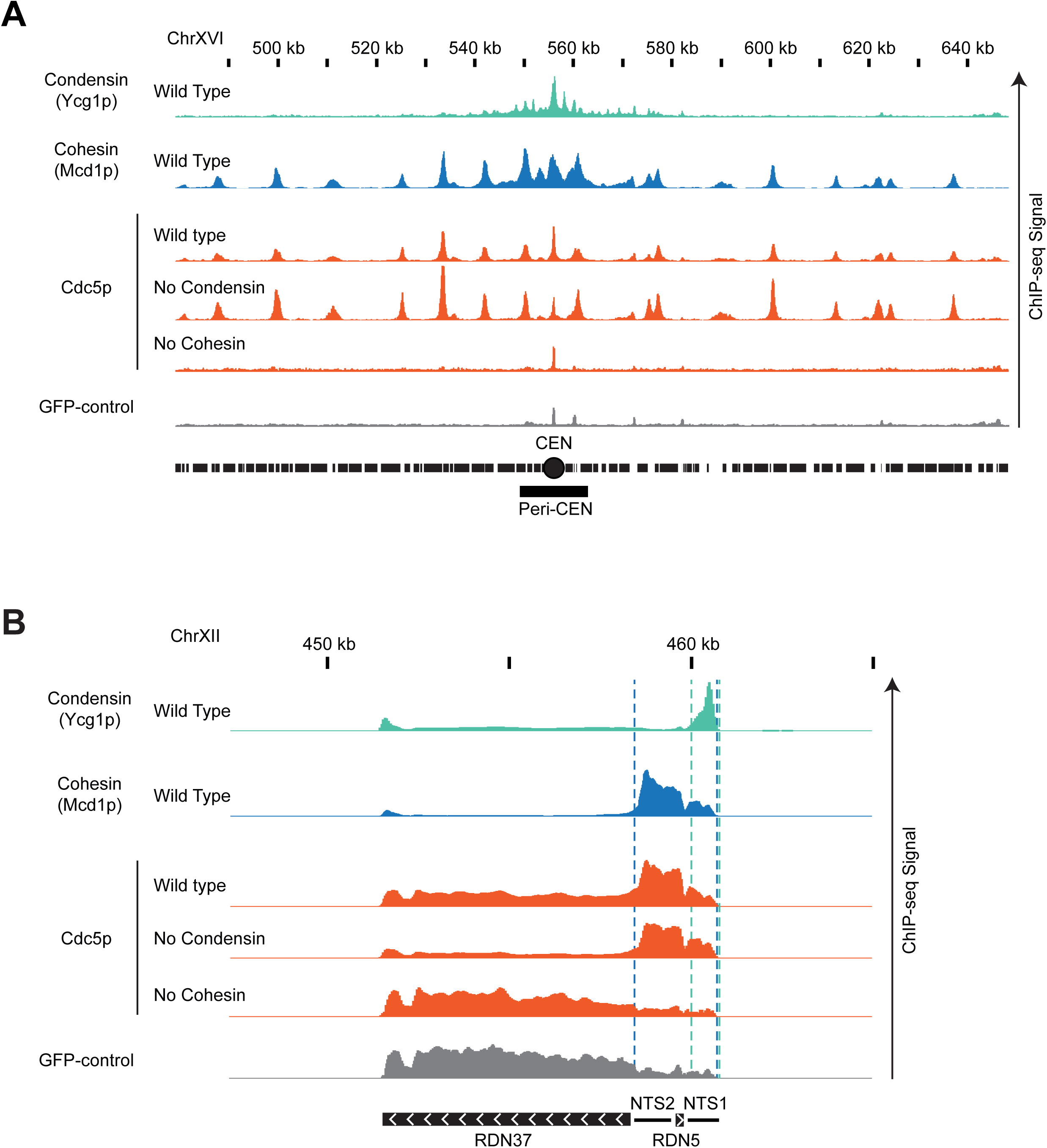
*Genome-wide localization of condensin, cohesin and Cdc5p.* **A.** Cohesin recruits Cdc5p at unique sequences genome-wide. ChIP-seq signals from a representative portion of chromosome XVI are presented with bar charts. The region presented includes the centromere (CEN black circle) and the pericentric region (peri-CEN black box). Signal coming from condensin subunit Ycg1p-Myc in wild type strain is in green, signal from cohesin subunit Mcd1p in wild type strain is in blue, signal coming from Cdc5p-Flag is in orange, and signal coming from GFP-control is in grey. Cdc5p ChIP-seq signal from wild type is in the first orange lane, from cells depleted of condensin subunit Brn1p-AID is in the second orange lane, from cells depleted of cohesin subunit Mcd1p-AID is in the third orange lane. All strains were depleted of the indicated AID-protein from G1 to mid-M and arrested in mid-M (staged mid-M). The scale is 0-15 for Ycg1p, 0-21 for Mcd1p, and 0-17 for Cdc5p. **B.** Cohesin recruits Cdc5p at the repetitive RDN locus. ChIP-seq signals from a portion of chromosome XII are presented with bar charts. ChIP-seq signal from strains as in (A). The region presented includes one copy of the rDNA repeat (9.1kb) made of the transcribed regions RND37 and RND5 (black box with white arrows indicating the direction of transcription) and the non-transcribed regions NTS1 and NTS2 (black line). Region of enrichment of cohesin signal is marked by two blue dashed lines (from NTS2 to NTS1). Region of enrichment of condensin signal is marked by two green dashed lines (end of NTS1). The scale is 0-5200 for Ycg1p, 0-2300 for Mcd1p, and 0-1700 for Cdc5p.

We then examined Brn1p, Mcd1p and Cdc5p binding patterns after depletion of Cdc5p-AID, Brn1p-AID or Mcd1p-AID. We depleted the relevant factor from G1 to mid-M using the staged mid-M regimen. The depleted factor could not be detected by western blot. The chromosomal binding pattern of Ycg1p-Myc did not change upon the depletion of either cohesin or Cdc5p (Figure 3—figure supplement 1; Figure 3—figure supplement 2; Figure 3—figure supplement 3). Similarly, the binding pattern of Mcd1p remained unchanged upon depletion of either condensin or Cdc5p (Figure 3—figure supplement 1; Figure 3—figure supplement 2; Figure 3—figure supplement 3). Thus, cohesin and condensin bind to specific sites on chromosomes independently of each other and Cdc5p. Therefore, cohesin and Cdc5p do not regulate RDN condensation by changing condensin localization to the RDN.

We then analyzed whether changes in the localization of Cdc5p might provide insights into RDN condensation. We examined cells in which Mcd1p-AID or Brn1p-AID were depleted from G1 to mid-M using the staged mid-M regimen. Upon Mcd1p-AID depletion, the binding of Cdc5p was abolished genome-wide (Figure 3; Figure 3—figure supplement 1; Figure 3—figure supplement 2; Figure 3—figure supplement 3), despite similar levels of total Cdc5p (Figure 3—figure supplement 4). This result recapitulated previous work showing the cohesin-dependent binding of Cdc5p at CARs (Mishra et al., 2016; Pakchuen et al., 2016). A few small peaks of binding were observed at centromere and RDN regions, but they coincided with false positives in the GFP-control.

In contrast, upon Brn1p-AID depletion, the pattern of Cdc5p was largely unaffected genome-wide (Figure 3; Figure 3—figure supplement 1; Figure 3—figure supplement 2; Figure 3— figure supplement 3). These results showed that the recruitment of Cdc5p to chromosomes (including the RDN) required the presence of cohesin over some period between G1 to mid-M

We reasoned the purpose of cohesin recruitment of Cdc5p was to promote localized Cdc5p binding and stimulation of condensin. Therefore, we predicted that a transient association of cohesin with the RDN would be sufficient to bring some Cdc5p to condensin. To test this possibility, we synchronized *MCD1-AID* cells in mid-M in the absence of auxin to allow cohesin, condensin, and Cdc5p to normally localize to the RDN. We then added auxin to deplete Mcd1p-AID and remove cohesin. We observed that Cdc5p was eliminated throughout the genome except at a portion of the condensin-bound regions of the RDN and centromeres (Figure 4; Figure 4—figure supplement 1). In these regions, we observed peaks of Cdc5p binding that were 3-5 fold greater than those observed when Mcd1p-AID was removed from G1 to mid-M. The remaining portion of the Cdc5p binding when Mcd1p-AID was depleted in mid-M, was not sufficient to promote proper condensation (Figure 4—figure supplement 2). To corroborate our ChIP-seq observations, we used ChIP followed by quantitative PCR (ChIP-qPCR) to follow the temporal dissociation of Cdc5p from the RDN at different time points after cohesin depletion in mid-M (Figure 4—figure supplement 3). We observed that Mcd1p binding disappeared from the RDN after 15 minutes with auxin. Consequently, Cdc5p binding also disappeared from the RDN, except at the site where condensin binds. Here, more than 50% of Cdc5p binding persisted, eventually dissipating to background level (90 minutes). Thus, as suggested from the ChIP-seq, transient Mcd1p binding to the RDN in mid-M is required for Cdc5p binding to the site of condensin binding. We propose that in wild type, high levels of cohesin at the RDN bring high levels of Cdc5p to neighboring condensin, promoting condensin activation and proper condensation.

**Figure 4:**
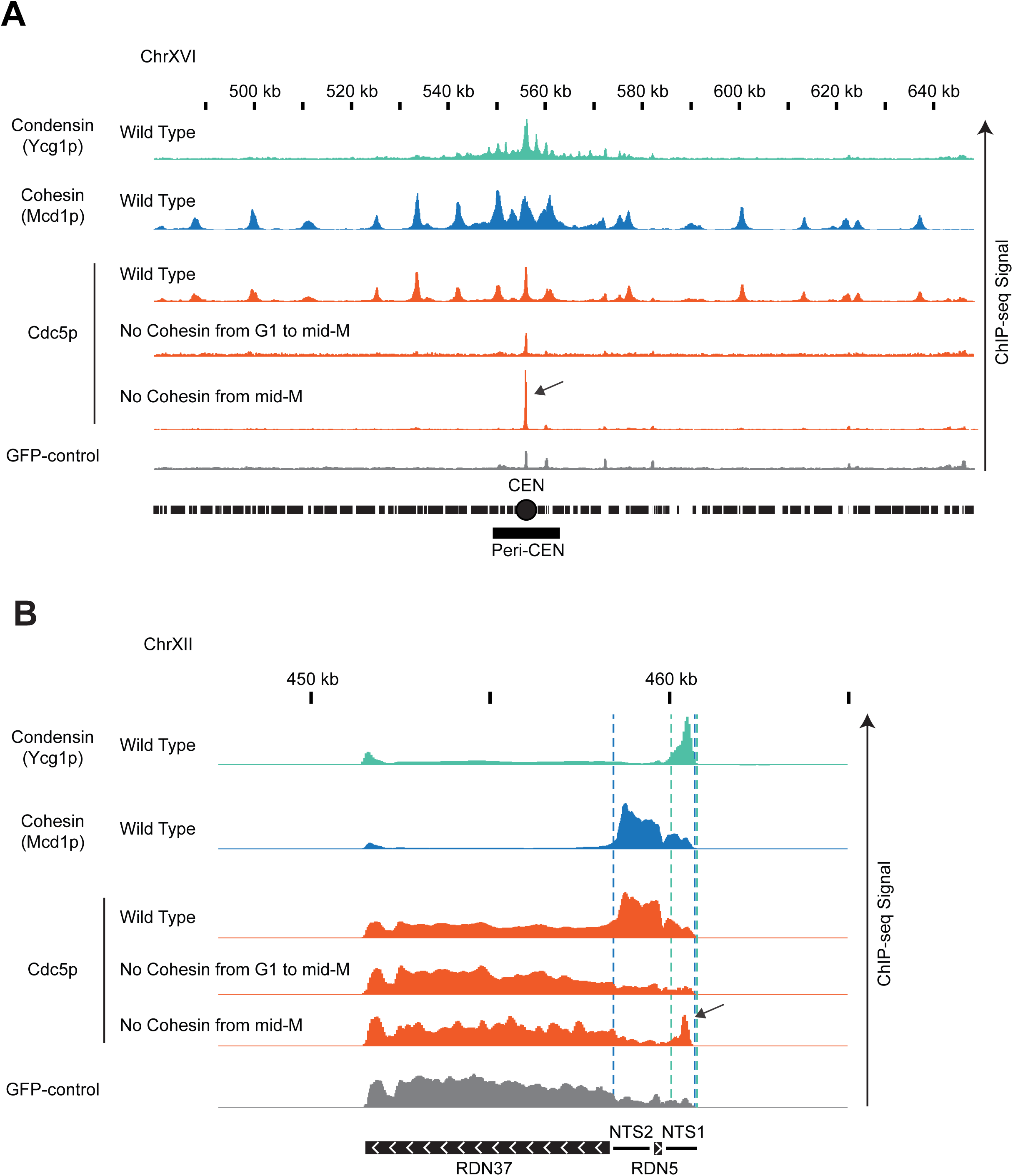
*Cdc5p is recruited by cohesin and brought to condensin at centromeres and rDNA.* **A.** Cdc5p is recruited by cohesin and brought to condensin at centromeres. ChIP-seq signals from a representative portion of chromosome XVI are presented with bar charts. The region presented includes the centromere (CEN black circle) and the pericentric region (peri-CEN black box). Signal coming from condensin subunit Ycg1p-Myc in wild type strain is in green, signal from cohesin subunit Mcd1p in wild type strain is in blue, signal coming from Cdc5p-V5 is in orange, and signal coming from GFP-control is in grey. Cdc5p ChIP-seq signal from wild type is in the first orange lane, from cells depleted of cohesin subunit Mcd1p-AID from G1 to mid-M is in the second orange lane, from cells depleted of cohesin subunit Mcd1p-AID in mid-M is in the third orange lane. Black arrow indicates enrichment of Cdc5p at the centromere when cohesin is transiently associated with chromosomes. The scale is 0-15 for Ycg1p, 0-21 for Mcd1p, and 0-17 for Cdc5p. **B.** Cdc5p is recruited by cohesin and brought to condensin at RDN. ChIP-seq signals from a portion of chromosome XII are presented with bar charts. ChIP-seq signal from strains as in (A). The region presented includes one copy of the rDNA repeat (9.1kb) made of the transcribed regions RND37 and RND5 (black box with white arrows indicating the direction of transcription) and the non-transcribed regions NTS1 and NTS2 (black line). Region of enrichment of cohesin signal is marked by two blue dashed lines (from NTS2 to NTS1). Region of enrichment of condensin signal is marked by two green dashed lines (end of NTS1). Black arrow indicates enrichment of Cdc5p at the condensin-bound region of the RDN when cohesin is transiently associated with chromosomes. The scale is 0-5200 for Ycg1p, 0-2300 for Mcd1p, and 0-1700 for Cdc5p.

### Localization of Cdc5p with CRISPR-Cas9 to the RDN partially suppresses the condensation defect of cohesin-depleted cells

If cohesin localization of Cdc5p to condensin at the RDN did indeed promote condensation, this function might be mimicked by ectopic localization of Cdc5p. To test this possibility, we engineered a strain harboring a fusion gene of Cdc5p and catalytically dead dCas9 under the control of the galactose promoter (Figure 5A). The fusion protein supported the growth of yeast as the sole source of Cdc5p (Figure 5—figure supplement 1A), indicating that this fusion protein retained activity. The strain with the fusion gene into *MCD1-AID* cells was transformed with plasmids containing either a guide RNA targeting the region of condensin binding at the RDN (NTS1 gRNA) or a distal guide RNA targeting 500 bp away (RDN5 gRNA) (Figure 5A). The successful target of the fusion to these two sites was corroborated by ChIP-qPCR (Figure 5—figure supplement 2A). These strains were subjected to the staged mid-M regimen to deplete Mcd1p-AID. The expression of dCas9-Cdc5p was induced from G1 by the addition of galactose to the media, and the ability of the targeted fusion to suppress RDN misfolding was assessed (Figure 5—figure supplement 1B).

**Figure 5:**
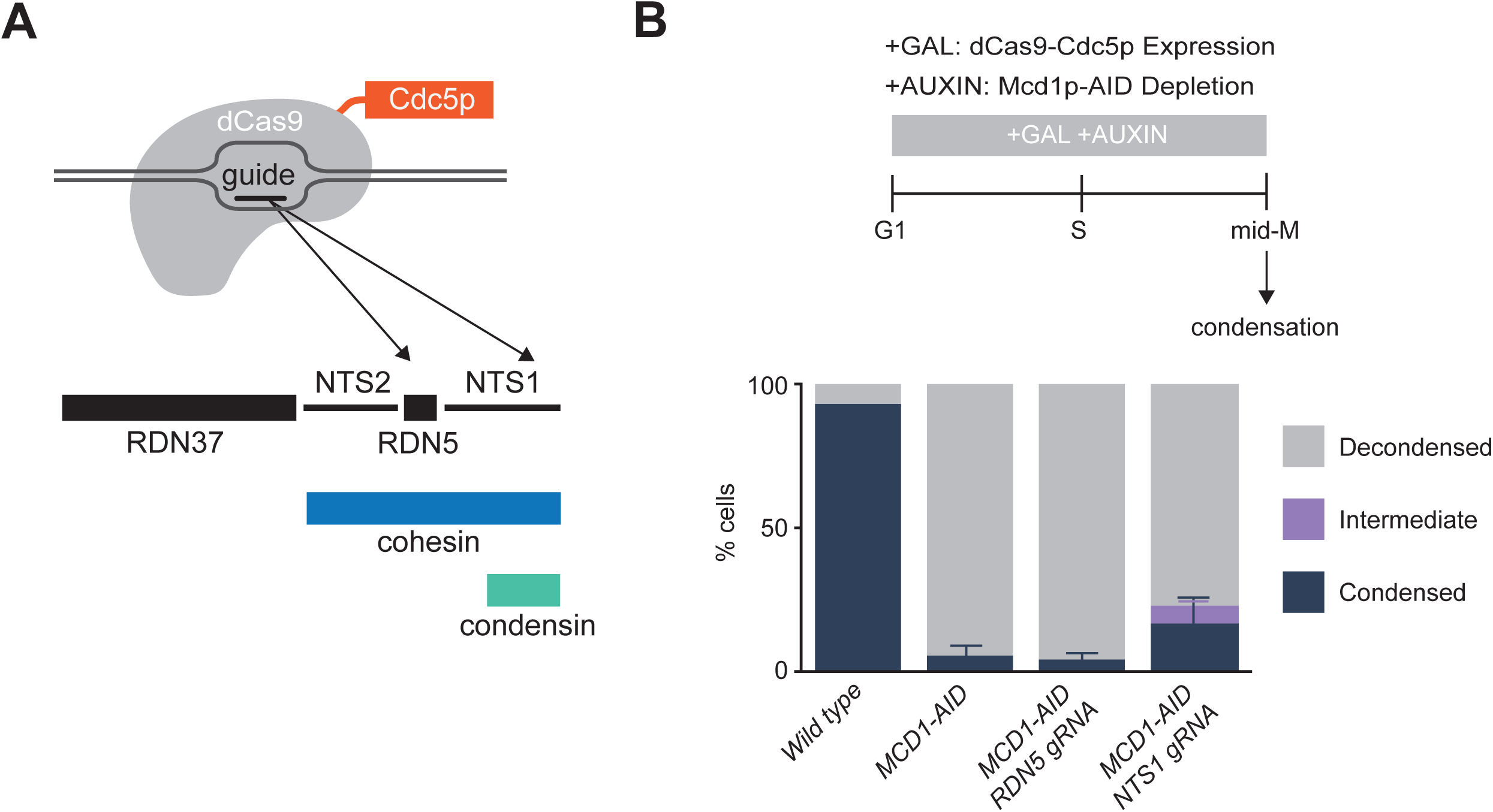
*Localization of Cdc5p with CRISPR-Cas9 to the condensin-bound RDN partially restores the RDN condensation defect of cohesin-depleted cells.* **A.** Galactose addition to the media drives the expression of a catalytically dead dCas9 fused to Cdc5p. Guide RNAs recruit dCas9-Cdc5p fusion protein either to the condensin-bound region NTS1 (RDN5 gRNA) or 500 bp to the left at RDN5 (RDN5 gRNA) where only cohesin is bound. **B.** *Top:* ‘Staged mid-M’ regimen used to assess condensation in mid-M. Cultures were synchronized in G1 and released in media containing galactose and auxin to drive the expression of dCas9-Cdc5p concomitantly to the depletion of Mcd1p-AID. Media also contained nocodazole to arrest cells in mid-M and assess condensation. *Bottom:* Wild type, Mcd1p depleted, Mcd1p depleted with dCas9-Cdc5p and RDN5 gRNA expressing, and Mcd1p depleted with dCas9-Cdc5p and NTS1 gRNA expressing cells were treated with ‘staged mid-M’ regimen and processed to score RDN condensation. Average of three experiments scoring 100 cells are shown and error bars represent the standard deviation.

In Mcd1p depleted cells, only 6% of the cells show normal condensed RDN loops (Figure 5B). In contrast, in the presence of guide RNA proximal to condensin, we observed 17% of cells with normally condensed loops, in addition to 6% partially condensed (Figure 5B; Figure 5—figure supplement 2B). The suppression of RDN misfolding could have been caused by the localization of Cdc5p immediately proximal to condensin, localization anywhere in the RDN, or by the excess of dCas9-Cdc5p fusion protein caused by galactose-induced overexpression. To distinguish between these possibilities, we examined RDN condensation when the overexpressed fusion was targeted 500 base pairs from condensin binding. No restoration of RDN condensation was observed, demonstrating that neither localization of the fusion anywhere in the RDN nor excess of the fusion protein was capable of promoting RDN condensation. Furthermore, the level of suppression of RDN misfolding by the NTS1-targeted fusion protein was underestimated by this assay. More than forty percent of the cells had lost the plasmid expressing the NTS1 gRNA prior to assessing condensation, generating false negatives for suppression. When we normalized to cells in the culture that retained the plasmid, the fraction of cells exhibiting partial or completely suppressed RDN folding was greater than 40%. These results show that ectopic targeting of Cdc5p specifically to sites of condensin binding in the RDN is capable of significantly replacing cohesin’s function in RDN condensation.

## Discussion

In this study, we have identified an interactome of early condensation factors (composed of cohesin, condensin, and the polo kinase Cdc5p) that control the establishment of chromosome condensation for the 1.4 megabases RDN locus of budding yeast. Our results suggest that cohesin recruits Cdc5p to the RDN and that Cdc5p actively promotes proper condensation, likely by stimulating condensin activity. When cohesin is absent, Cdc5p stimulates condensin to misfold chromosomes into an irreversibly decondensed state. Our data suggest that productive condensin activity is reliant on cohesin’s spatial regulation of Cdc5p. Altogether, our study describes a novel mechanism by which cohesin regulates condensin activity and begins to provide mechanistic insight into how SMC complexes can be spatially and temporally coordinated to achieve their functions.

### Cdc5p promotes the establishment and maintenance of condensation prior to anaphase

In this study, we corroborate a previous study that showed that condensation of the RDN in mid-M prior to anaphase requires Cdc5p (polo) kinase as well as CDK kinase (Walters et al., 2014). We show that Cdc5p kinase activity is required for both the establishment and maintenance of RDN condensation in mid-M of budding yeast, likely by modulating condensin. We propose that Cdc5p-dependent phosphorylation of condensin or a condensin regulator is the key step to establish condensation, and that continued phosphorylation is necessary to maintain condensation. We favor condensin as the likely key substrate since a previous study showed that Cdc5p phosphorylates condensin prior to anaphase (St-Pierre et al., 2009). The requirement for CDK and Cdc5p kinases in mid-M RDN condensation might reflect that the binding of polo kinases to their substrates often requires their prior phosphorylation by CDK. Alternatively, CDKs and Cdc5p may regulate different condensin activities like tethering DNA, supercoiling DNA and extruding DNA loops (Ganji et al., 2018; Hagstrom et al., 2002; Kimura et al., 2001; Kimura and Hirano, 1997; Lavoie et al., 2002; St-Pierre et al., 2009). Consistent with this idea, CDK phosphorylation is limited to the Smc4p subunit of condensin (Robellet et al., 2015) while Cdc5p phosphorylates Ycg1p, Ycs4p, and Brn1p subunits (St-Pierre et al., 2009).

### Cdc5 recruitment to the condensin-bound region of the RDN modulates condensin to promote proper RDN condensation

In this study, we provide important clues on how the regulation of condensin by cohesin and Cdc5p is important for the folding of the RDN. First, cohesin is required for the initial recruitment of Cdc5p to chromosomes in mid-M (this study; Mishra et al., 2016; Pakchuen et al., 2016). We show that cohesin recruitment of Cdc5p to chromosomes promotes Cdc5p binding to neighboring condensin. We suggest that cohesin recruitment either directly hands off Cdc5p to neighboring condensin or generates a high local concentration of Cdc5p that promotes its efficient interaction with condensin. By constraining the interaction of Cdc5p with condensin at the RDN, condensin activity is spatially controlled to promote proper RDN folding. In support of this model, we show that cohesin’s recruitment function can be replaced by ectopic localization of Cdc5p to the RDN by its fusion with Cas9.

The spatial activation of condensin through a localized kinase adds a critical second feature to the regulation of condensation. Indeed, modulating a kinase’s recruitment to its target protein is a very common theme in kinase regulation (Ferrell and Cimprich, 2003; Langeberg and Scott, 2015). It will be interesting to determine whether the regulation of condensin by CDKs and aurora kinase are also controlled spatially. However, the fact that condensation was not fully restored by the Cas9-Cdc5p fusion suggests that proper Cdc5p phosphorylation of condensin involves more than simply its general localization to the RDN bound condensin. Both the number of Cdc5p at the RDN and its exact spatial orientation relative to condensin likely differ between the fusion and cohesin bound Cdc5p. Alternatively, cohesin may have a Cdc5p-independent role in condensation. Discovering these features of condensin regulation will require additional studies.

### Cohesin localized Cdc5p and RDN misfolding

Cells lacking cohesin have three phenotypes that inform on the potential molecular function for cohesin-dependent localization of Cdc5p to the RDN. First, the RDN is irreversibly misfolded into a disorganized state (this study; Lavoie et al., 2002). Second, this RDN misfolding is driven by condensin and Cdc5p (this study; Lavoie et al., 2002). Third, Cdc5p phosphorylates Ycg1p subunit of condensin with a different phospho-pattern (this study). To explain these observations, we suggest that the complex pattern of Cdc5p phosphorylation of condensin is to coordinate condensin’s different activities. In wild type cells, cohesin recruits Cdc5p to condensin, mediating a temporally and spatially controlled Cdc5p phosphorylation of condensin. This regulation of condensin phosphorylation correctly activates the different activities of condensin, resulting in a properly folded RDN. In the absence of cohesin, the delocalized Cdc5p still phosphorylates condensin, but with a different phosphorylation pattern, leading to the uncoordinated control of its activities and RDN misfolding. Inactivating condensin or Cdc5p prevents the formation of the uncoordinated condensin, eliminating RDN misfolding. Identifying these key cohesin-dependent phosphorylation sites will require extensive structural and genetic assays.

In summary, the co-localization of cohesin, polo kinase and condensin underlies a regulatory pathway that spatially modulates condensin function to generate a thin loop of condensed RDN. The pericentric regions are the only other genomic loci in yeast showing similar co-localization of cohesin, condensin, and polo kinase. We predict that these factors regulate condensation of the pericentric regions to generate thinly condensed structures as they do at the RDN. In yeast, these structures would be difficult to detect cytologically because of the small size of the pericentric regions (∼10 kilobases). Intriguingly, the pericentric regions are much larger in animals and plants. These regions are visible as thinner areas of the mitotic chromosomes also known as primary constrictions. Constrictions are also observed in the nucleolus organizer regions, the rDNA loci of multicellular organisms. We propose that these constrictions are the result of the same regulatory pathway we described here for the condensation of yeast RDN. Indeed, in vitro condensed chromosomes without cohesin and polo kinase lack constrictions (Shintomi et al., 2015). In vivo, cohesin and polo kinase depleted mammalian cells also fail to form primary constrictions (Gandhi et al., 2006). More broadly this regulatory pathway may serve as a general paradigm for how cohesin, condensin, and other SMC complexes communicate to achieve proper chromosome structure in other biological processes.

---

**Figure 1—figure supplement 1:**
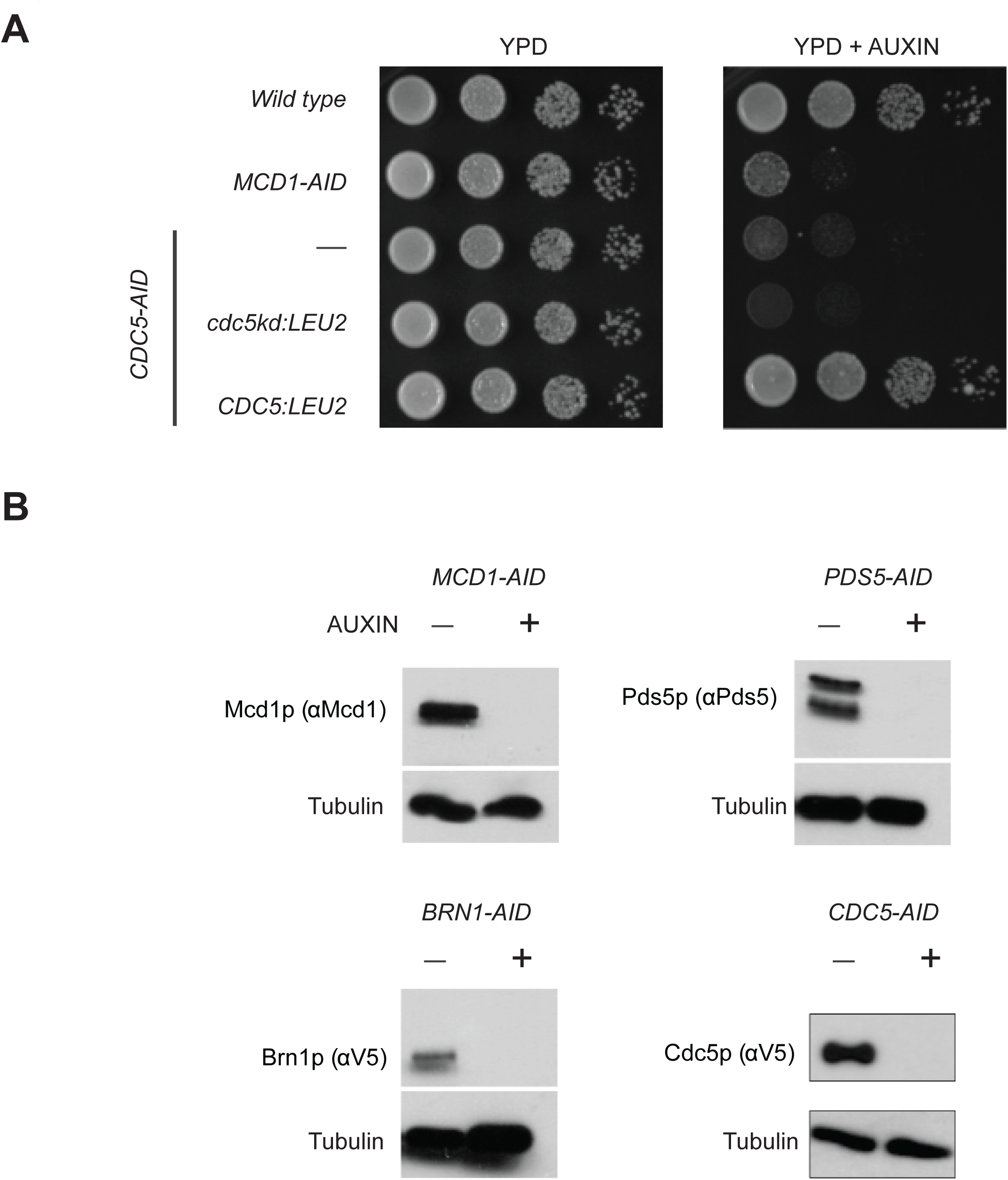
*Characterization of AID-tagged strains.* **A.** Cdc5p depletion using the AID degron system reduces viability. Asynchronous cells were serially diluted and plated on YPD (left) or YPD with 750uM auxin (right). Plates were then incubated at 23C for three days and imaged. Wild Type, Mcd1p-AID, Cdc5p-AID, Cdc5-AID with expression of Cdc5p kinase-dead from the LEU2 locus, and Cdc5p-AID with expression of Cdc5p from the LEU2 locus cells were plated. **B.** Depletion of AID-tagged proteins assayed by Western blotting. *CDC5-AID, BRN1-AID, MCD1-AID, PDS5-AID,* strains were grown to mid-log in YPD. Cells were then split and grown either in the presence or absence of auxin for 1h. Cells were then pelleted and protein extracted by TCA as described in the materials and methods. Western blot analysis used antibodies for the endogenous Mcd1p and Pds5p, anti V5 for Brn1p-3V5 and Cdc5p-3V5. Tubulin antibody was used for loading control

**Figure 1—figure supplement 2:**
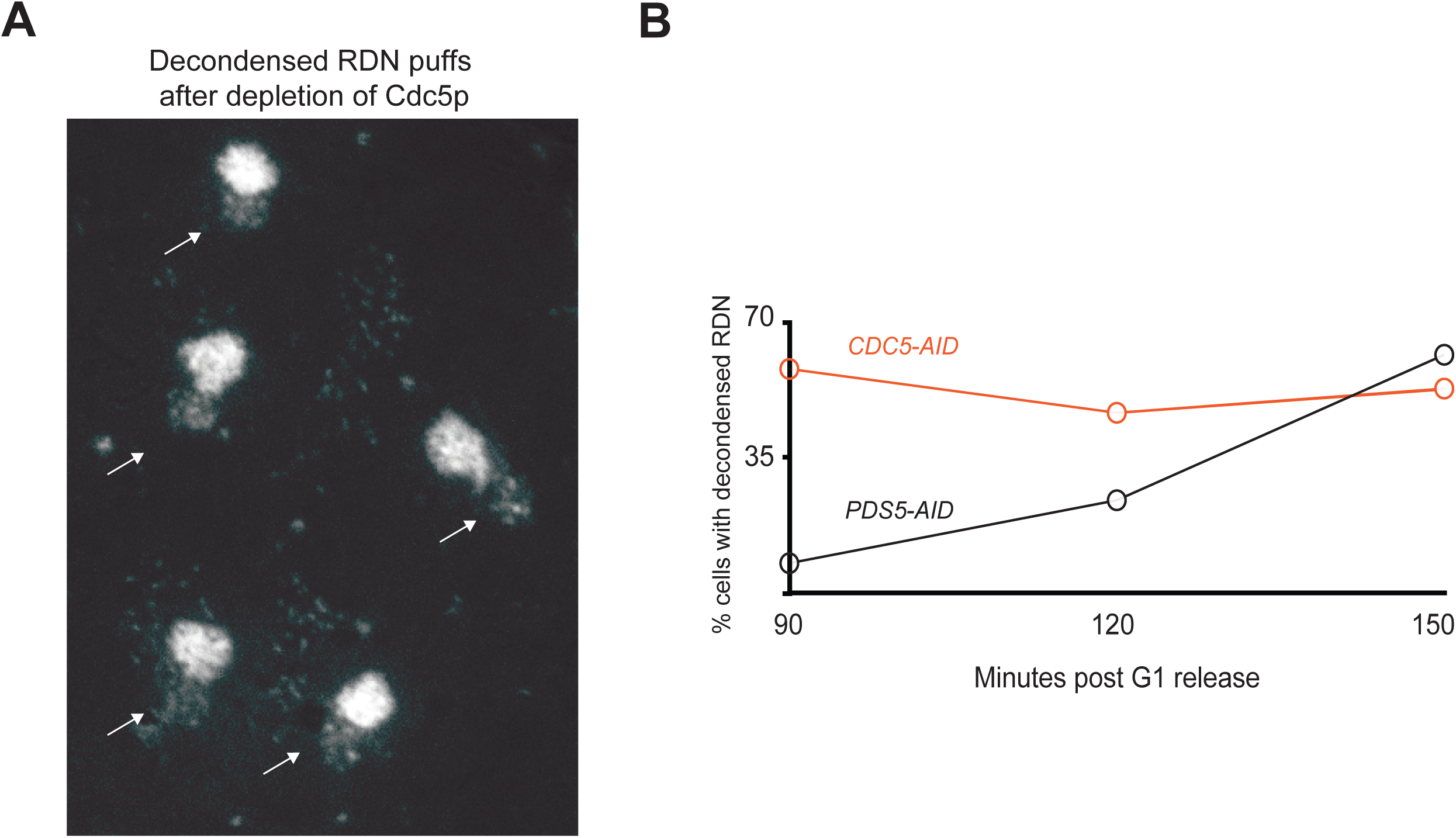
*Cdc5p depletion results in loss of RDN condensation* **A.** Loss of rDNA condensation after Cdc5p-AID depletion. Representative image showing chromosome spreads of Cdc5p-AID cells after auxin depletion. The puffed RDN next to the bulk of the chromosomes indicates loss of condensation in the RDN. **B.** Cdc5p is required to establish condensation of the RDN. Pds5p depleted and Cdc5p depleted, cells were treated with ‘‘Staged mid-M’ regimen used to assess strains function in mid-M. Cultures were synchronized in G1 and released in media containing auxin and nocodazole, to depleted AID-tagged protein and arrest cells in mid-M. RDN condensation assayed at 30mn intervals after release from G1 to mid-M. 90 minutes is the earliest time point were RDN condensation is clearly measurable. The average percentage of cells with decondensed RDN (100 cells) are reported.

**Figure 1—figure supplement 3:**
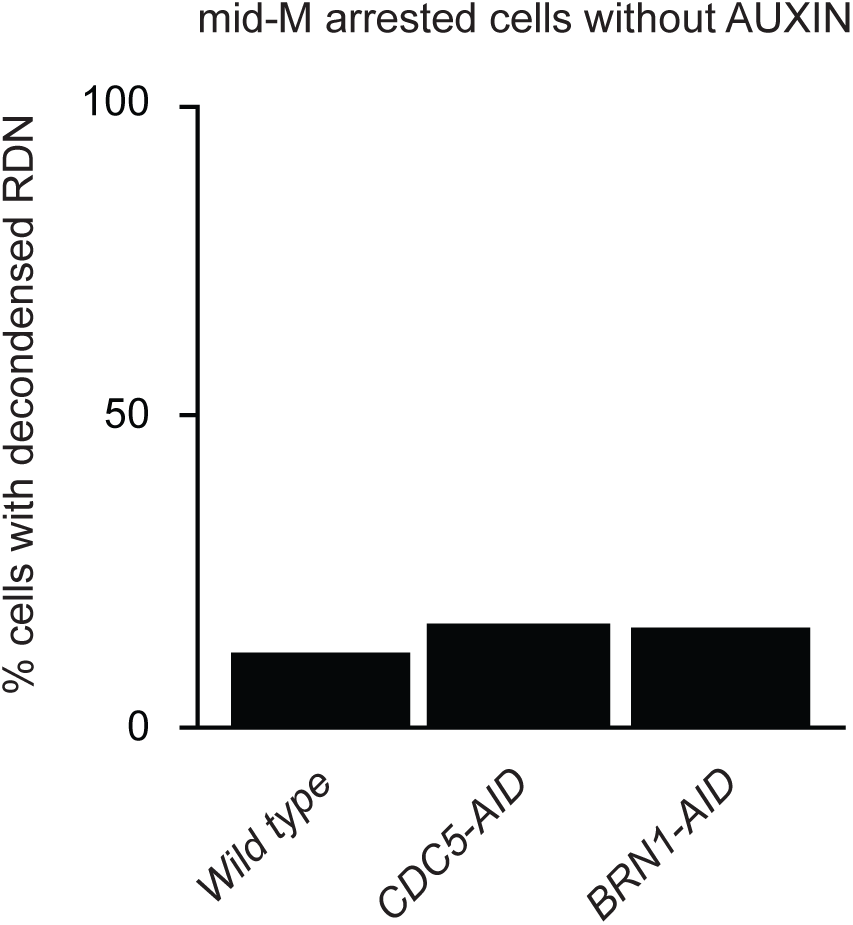
*Addition of the AID-tag to Cdc5p and Brn1 does not affect RDN condensation in the absence of auxin* rDNA condensation levels of AID-tagged strains are comparable to wild type without auxin. Wild type, Cdc5p depleted, and Brn1p depleted cells were treated as in B omitting auxin addition and processed to make chromosome spreads to score RDN condensation. 200 cells were scored.

**Figure 2—figure supplement 1:**
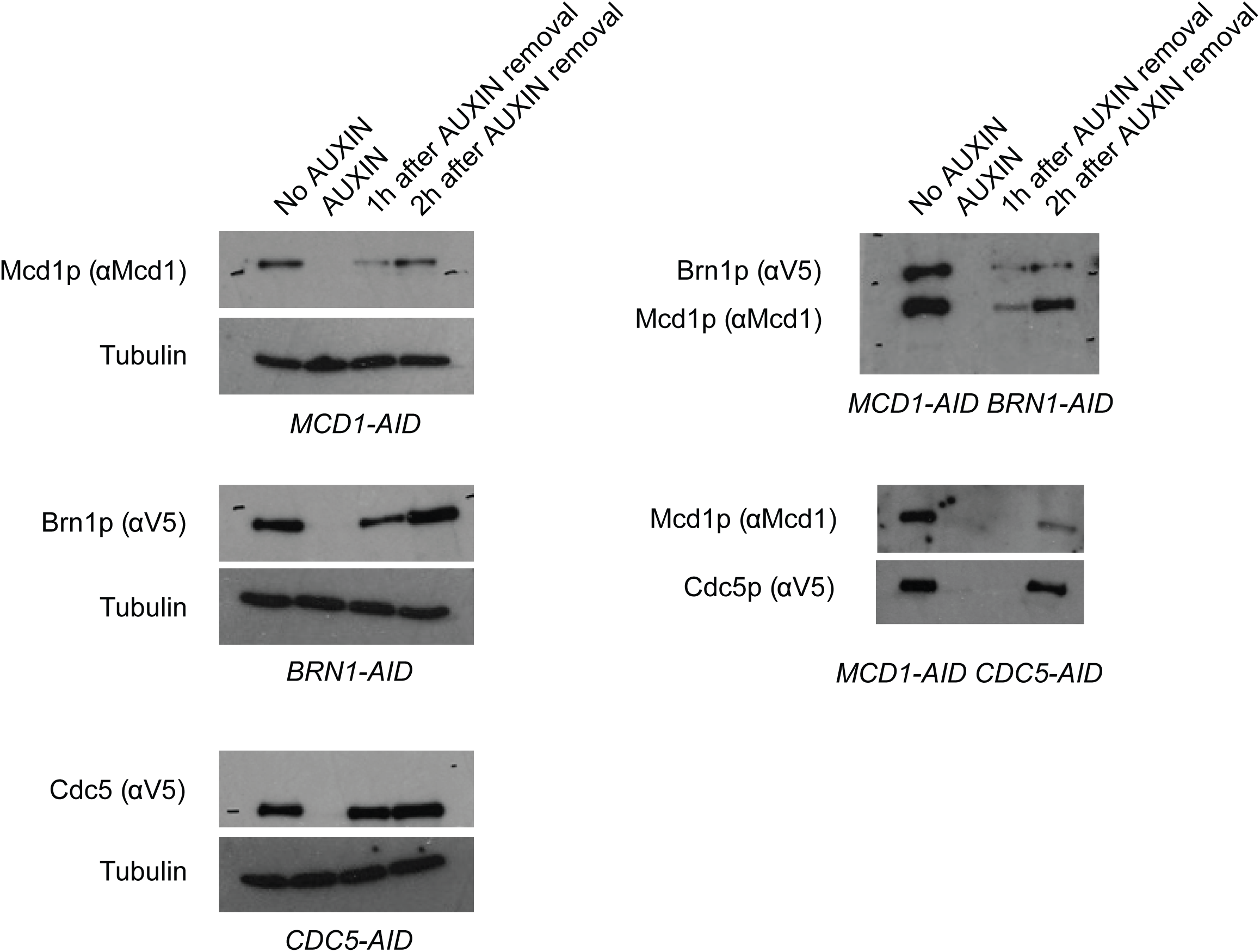
*AID-tagged protein levels analyzed after depletion and re-expression in the ‘add later’ experiment.* Depletion followed by expression of AID-tagged proteins assayed by Western blotting. Strains were subjected to the ‘Add later’ regimen. Indicated cultures were arrested in G1 (>95% schmooed morphology), and AID-tagged proteins (Mcd1p-AID, Brn1p-AID, Cdc5p-AID, MCD1-AID/BRN1-AID, and MCD1-AID/CDC5-AID) were depleted by the addition of auxin. G1 arrest was relieved by resuspending in media with auxin and nocodazole to arrest cells in mid-M. Cells were then resuspended in media containing only nocodazole for two hours to allow the expression of the previously depleted AID-tagged proteins. Cells were then pelleted and protein extracted by TCA as described in the materials and methods. Western blot analysis used antibodies for the endogenous Mcd1p, anti V5 for Brn1p-3V5 and Cdc5p-3V5. Tubulin antibody was used for loading control.

**Figure 3—figure supplement 1:**
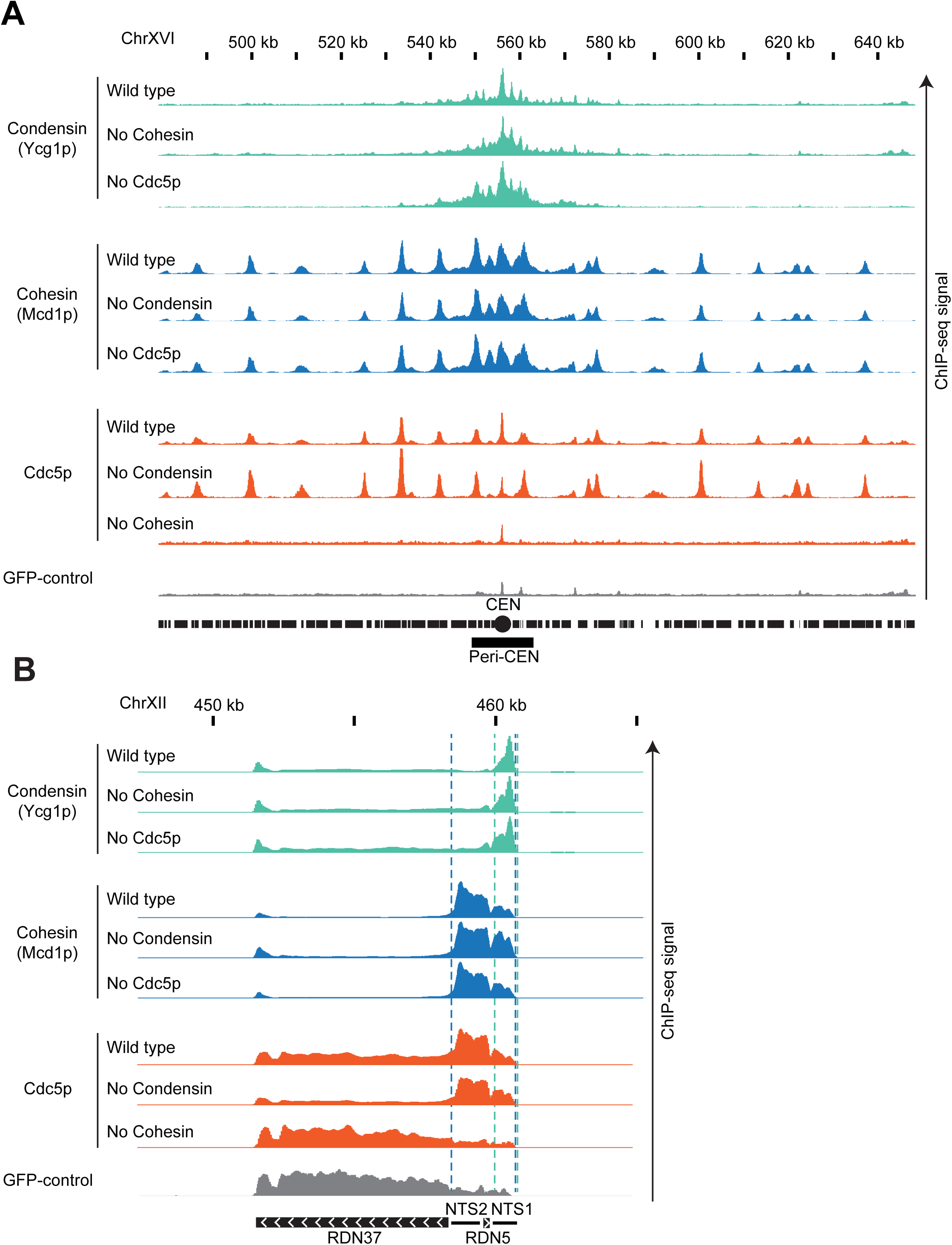
*Cohesin and condensin bind to specific sites on chromosomes independently of each other and Cdc5p.* **A.** Cohesin, condensin and Cdc5p localization at unique sequences genome-wide. ChIP-seq signals from a representative portion of chromosome XVI are presented with bar charts. The region presented includes the centromere (CEN black circle) and the pericentric region (peri-CEN black box). Signal coming from condensin subunit Ycg1p-Myc in wild type strain is in green, signal from cohesin subunit Mcd1p in wild type strain is in blue, signal coming from Cdc5p-Flag is in orange, and signal coming from GFP-control is in grey. Ycg1p-Myc signal from wild type is in the first green lane, from cells depleted of cohesin subunit Mcd1p-AID is in the second green lane, from cells depleted of Cdc5p-AID is in the third green lane. Mcd1p signal from wild type is in the first blue lane, from cells depleted of condensin subunit Brn1p-AID is in the second blue lane, from cells depleted of Cdc5p-AID is in the third blue lane.Cdc5p ChIP-seq signal from wild type is in the first orange lane, from cells depleted of condensin subunit Brn1p-AID is in the second orange lane, from cells depleted of cohesin subunit Mcd1p-AID is in the third orange lane. All strains were depleted of the indicated AID-protein from G1 to mid-M and arrested in mid-M (staged mid-M). The scale is 0-15 for Ycg1p, 0-21 for Mcd1p, and 0-17 for Cdc5p. **B.** Cohesin, condensin and Cdc5p localization at the repetitive RDN locus. ChIP-seq signals from a portion of chromosome XII are presented with bar charts. ChIP-seq signal from strains as in (A). The region presented includes one copy of the rDNA repeat (9.1kb) made of the transcribed regions RND37 and RND5 (black box with white arrows indicating the direction of transcription) and the non-transcribed regions NTS1 and NTS2 (black line). Region of enrichment of cohesin signal is marked by two blue dashed lines (from NTS2 to NTS1). Region of enrichment of condensin signal is marked by two green dashed lines (end of NTS1). The scale is 0-5200 for Ycg1p, 0-2300 for Mcd1p, and 0-1700 for Cdc5p.

**Figure 3—figure supplement 2:**
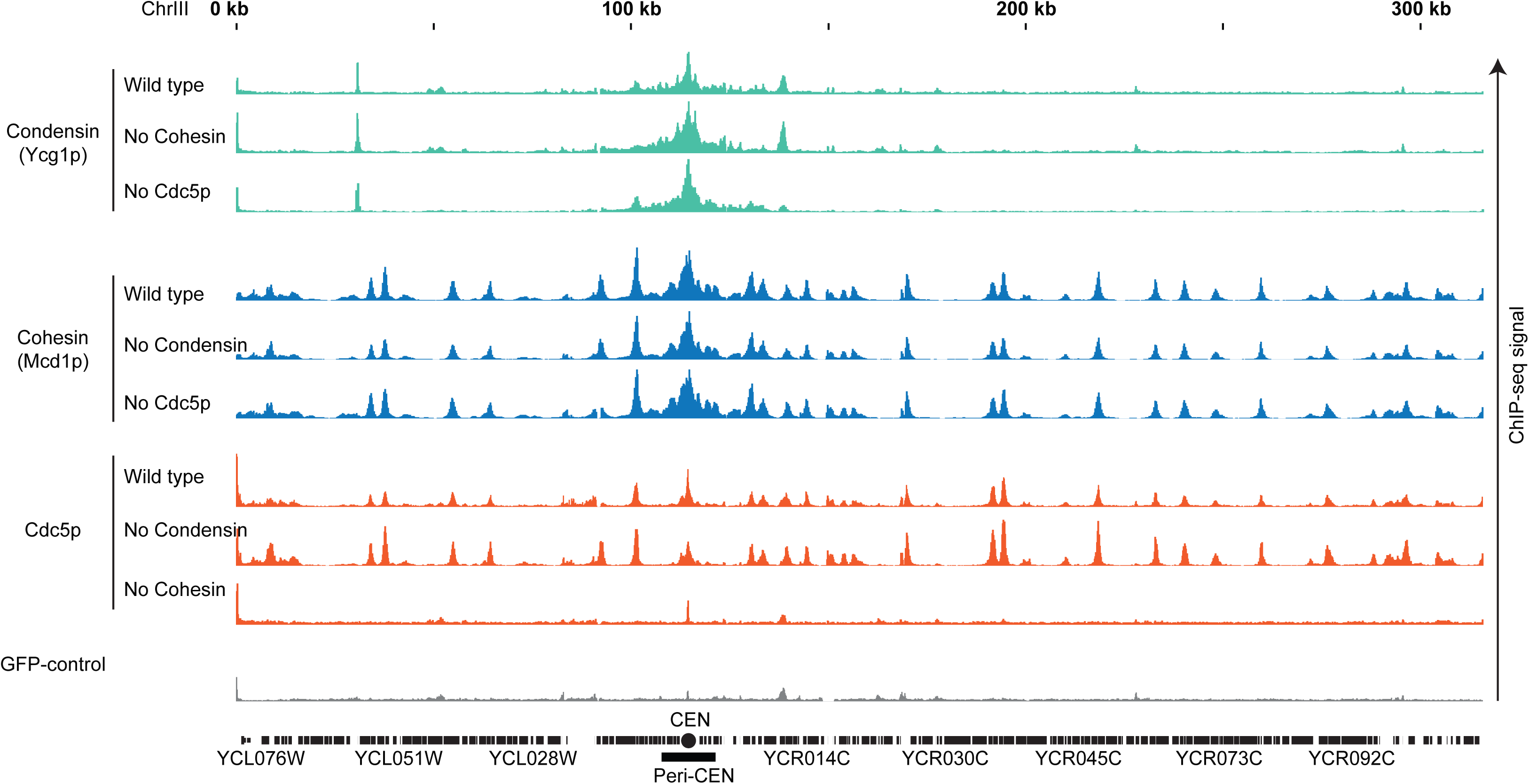
*Cohesin, condensin, and Cdc5p bind to specific sites on chromosome III.* Cohesin, condensin and Cdc5p localization at unique sequences genome-wide. ChIP-seq signals from a representative portion of chromosome III are presented with bar charts. The region presented includes the centromere (CEN black circle) and the pericentric region (peri-CEN black box). Signal coming from condensin subunit Ycg1p-Myc in wild type strain is in green, signal from cohesin subunit Mcd1p in wild type strain is in blue, signal coming from Cdc5p-Flag is in orange, and signal coming from GFP-control is in grey. Ycg1p-Myc signal from wild type is in the first green lane, from cells depleted of cohesin subunit Mcd1p-AID is in the second green lane, from cells depleted of Cdc5p-AID is in the third green lane. Mcd1p signal from wild type is in the first blue lane, from cells depleted of condensin subunit Brn1p-AID is in the second blue lane, from cells depleted of Cdc5p-AID is in the third blue lane.Cdc5p ChIP-seq signal from wild type is in the first orange lane, from cells depleted of condensin subunit Brn1p-AID is in the second orange lane, from cells depleted of cohesin subunit Mcd1p-AID is in the third orange lane. All strains were depleted of the indicated AID-protein from G1 to mid-M and arrested in mid-M (staged mid-M). The scale is 0-15 for Ycg1p, 0-21 for Mcd1p, and 0-17 for Cdc5p.

**Figure 3—figure supplement 3:**
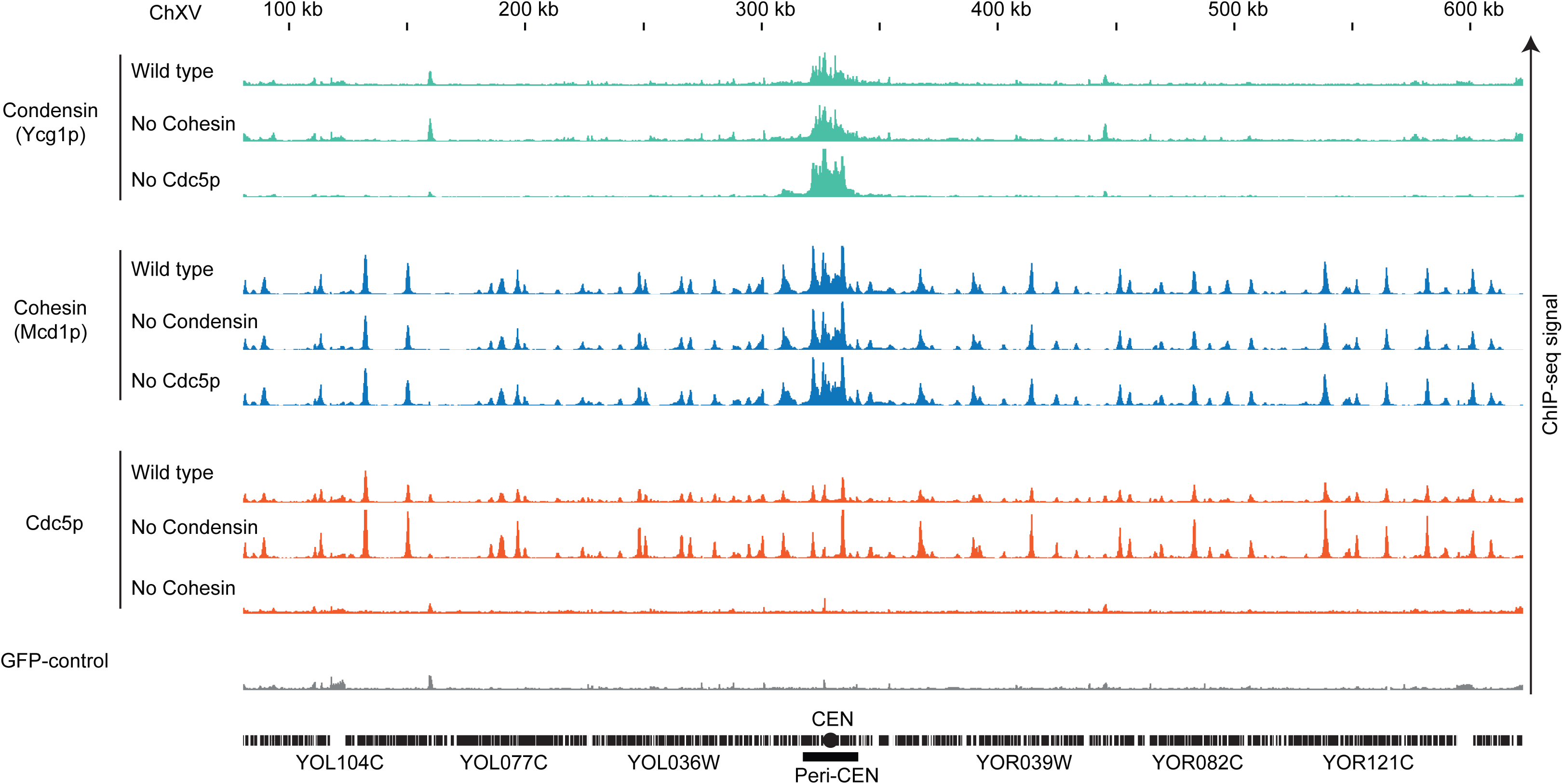
*Cohesin, condensin, and Cdc5p bind to specific sites on chromosome XV.* Cohesin, condensin and Cdc5p localization at unique sequences genome-wide. ChIP-seq signals from a representative portion of chromosome XV are presented with bar charts. The region presented includes the centromere (CEN black circle) and the pericentric region (peri-CEN black box). Signal coming from condensin subunit Ycg1p-Myc in wild type strain is in green, signal from cohesin subunit Mcd1p in wild type strain is in blue, signal coming from Cdc5p-Flag is in orange, and signal coming from GFP-control is in grey. Ycg1p-Myc signal from wild type is in the first green lane, from cells depleted of cohesin subunit Mcd1p-AID is in the second green lane, from cells depleted of Cdc5p-AID is in the third green lane. Mcd1p signal from wild type is in the first blue lane, from cells depleted of condensin subunit Brn1p-AID is in the second blue lane, from cells depleted of Cdc5p-AID is in the third blue lane.Cdc5p ChIP-seq signal from wild type is in the first orange lane, from cells depleted of condensin subunit Brn1p-AID is in the second orange lane, from cells depleted of cohesin subunit Mcd1p-AID is in the third orange lane. All strains were depleted of the indicated AID-protein from G1 to mid-M and arrested in mid-M (staged mid-M). The scale is 0-15 for Ycg1p, 0-21 for Mcd1p, and 0-17 for Cdc5p.

**Figure 3—figure supplement 4:**
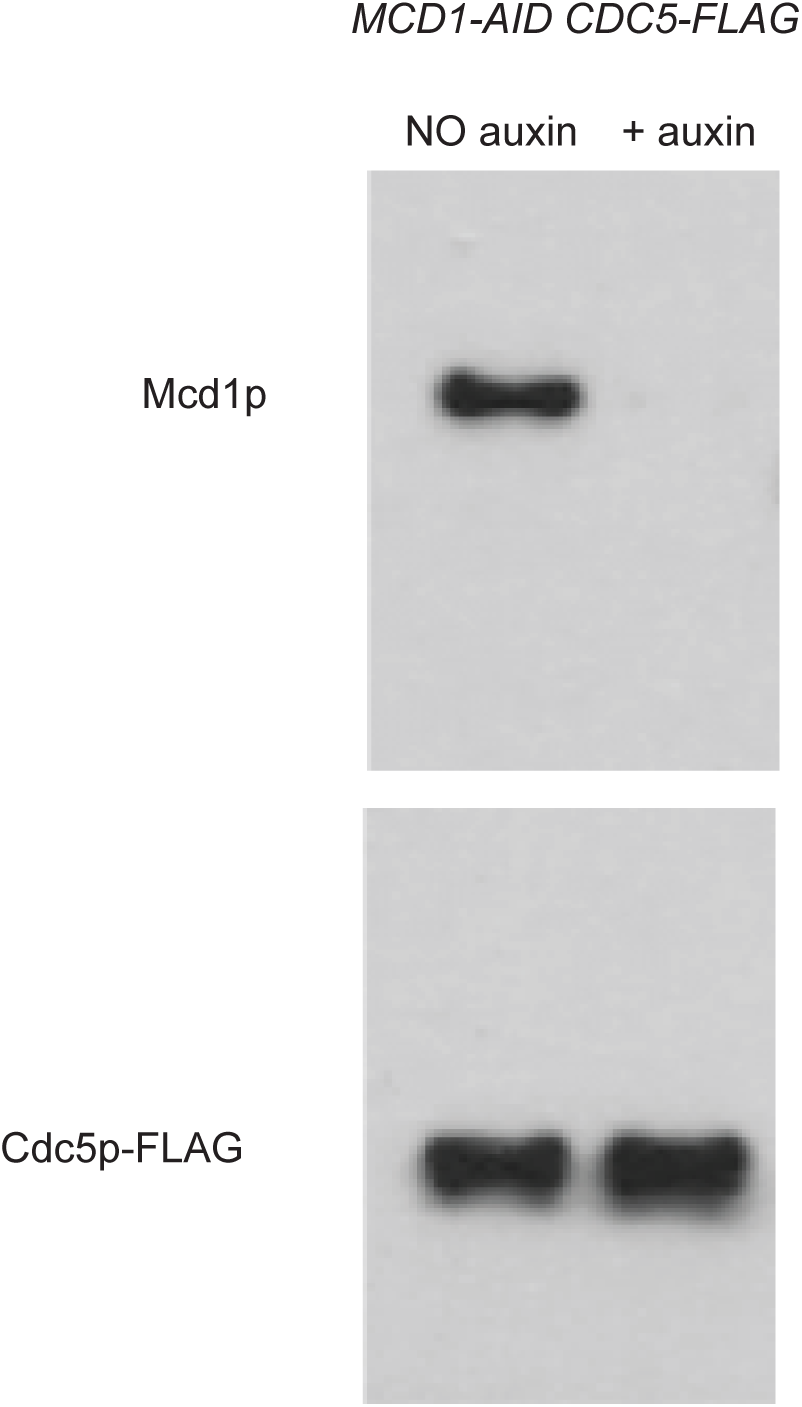
*Cdc5p levels after Mcd1p depletion analyzed by western blotting.* Depletion of Mcd1p-AID does not affect levels of Cdc5p. Indicated cultures were arrested in G1 (>95% schmooed morphology), and Mcd1p-AID was depleted by the addition of auxin. G1 arrest was relieved by resuspending in media with auxin and nocodazole to arrest cells in mid-M. Cells were then pelleted and protein extracted by TCA as described in the materials and methods. Western blot analysis used antibodies for the endogenous Mcd1p, anti FLAG for Cdc5p-FLAG.

**Figure 4—figure supplement 1:**
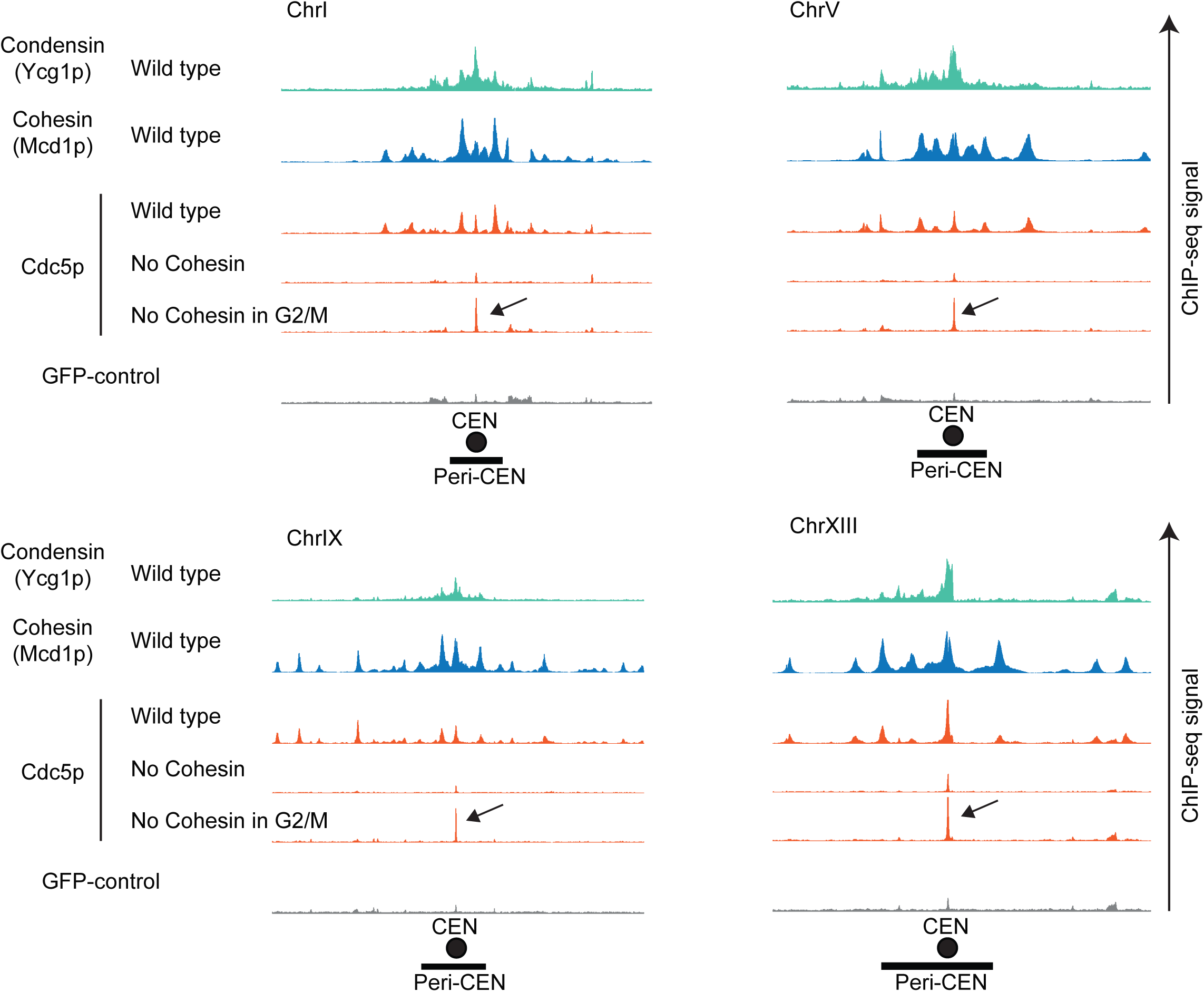
*Cdc5p binding to centromeres is revealed by a transient association of Mcd1p with chromosomes.* Cdc5p is recruited by cohesin and brought to condensin at centromeres. ChIP-seq signals from a representative portion of chromosomes are presented with bar charts. The region presented includes the centromere (CEN black circle) and the pericentric region (peri-CEN black box). Signal coming from condensin subunit Ycg1p-Myc in wild type strain is in green, signal from cohesin subunit Mcd1p in wild type strain is in blue, signal coming from Cdc5p-V5 is in orange, and signal coming from GFP-control is in grey. Cdc5p ChIP-seq signal from wild type is in the first orange lane, from cells depleted of cohesin subunit Mcd1p-AID from G1 to mid-M is in the second orange lane, from cells depleted of cohesin subunit Mcd1p-AID in mid-M is in the third orange lane. Black arrow indicates enrichment of Cdc5p at the centromere when cohesin is transiently associated with chromosomes. The scale is 0-15 for Ycg1p, 0-21 for Mcd1p, and 0-17 for Cdc5p.

**Figure 4—figure supplement 2:**
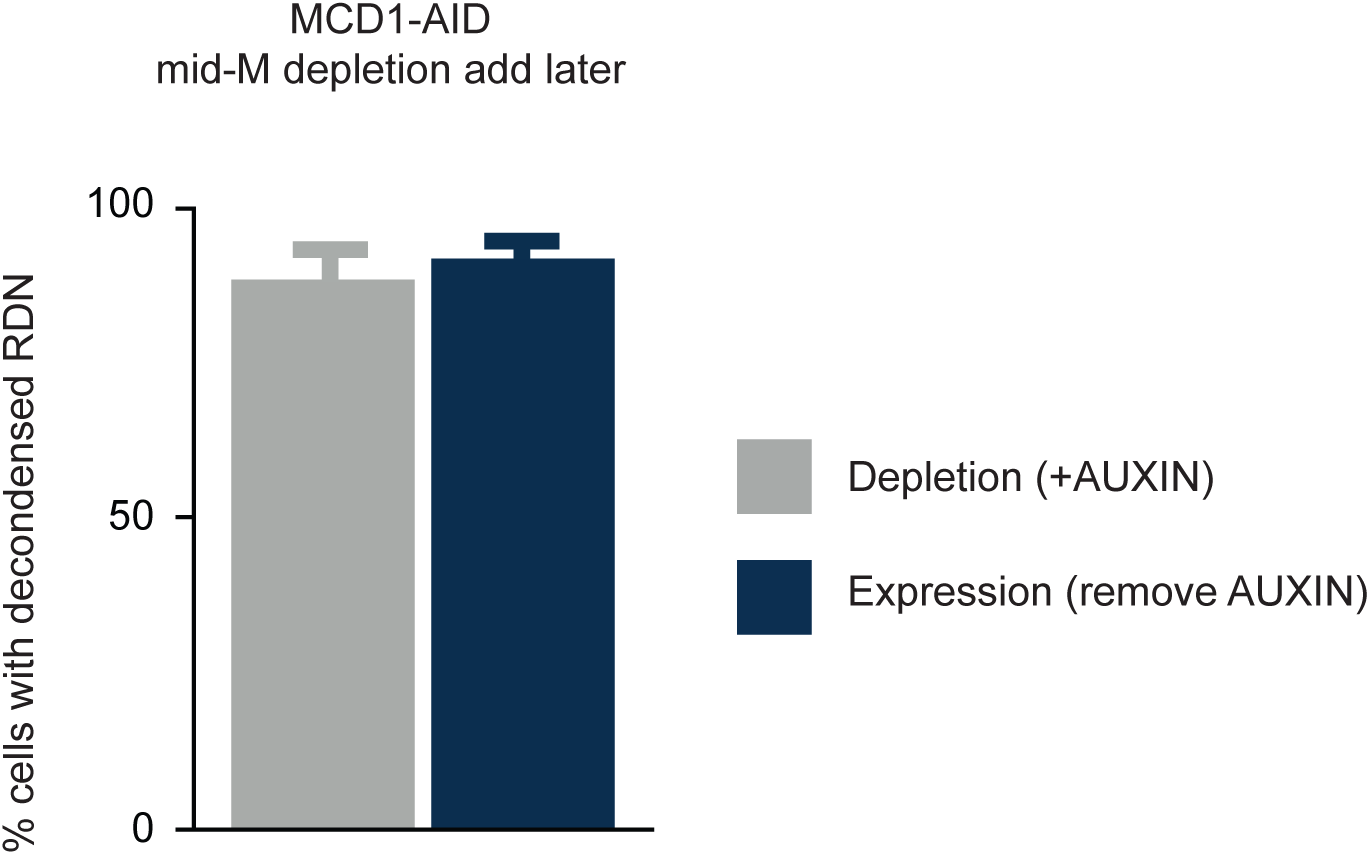
*Mcd1p transient association with chromosomes does not restore condensation of the RDN in Mcd1p depleted cells.* Depletion of Cdc5p results in loss of maintenance of condensation. Mcd1p-AID cells were arrested in mid-M using nocodazole. The arrest was confirmed by analysis of bud morphology (>95% large budded cells). Auxin was added to ensure Mcd1p-AID degradation for 90min (depletion grey bar). Mcd1p-AID was then expressed by washing out auxin (expression blue bar). Cells were processed to make chromosome spreads to score RDN condensation. Two independent experiments with 200 cells were scored and error bars indicate the standard deviation.

**Figure 4—figure supplement 3:**
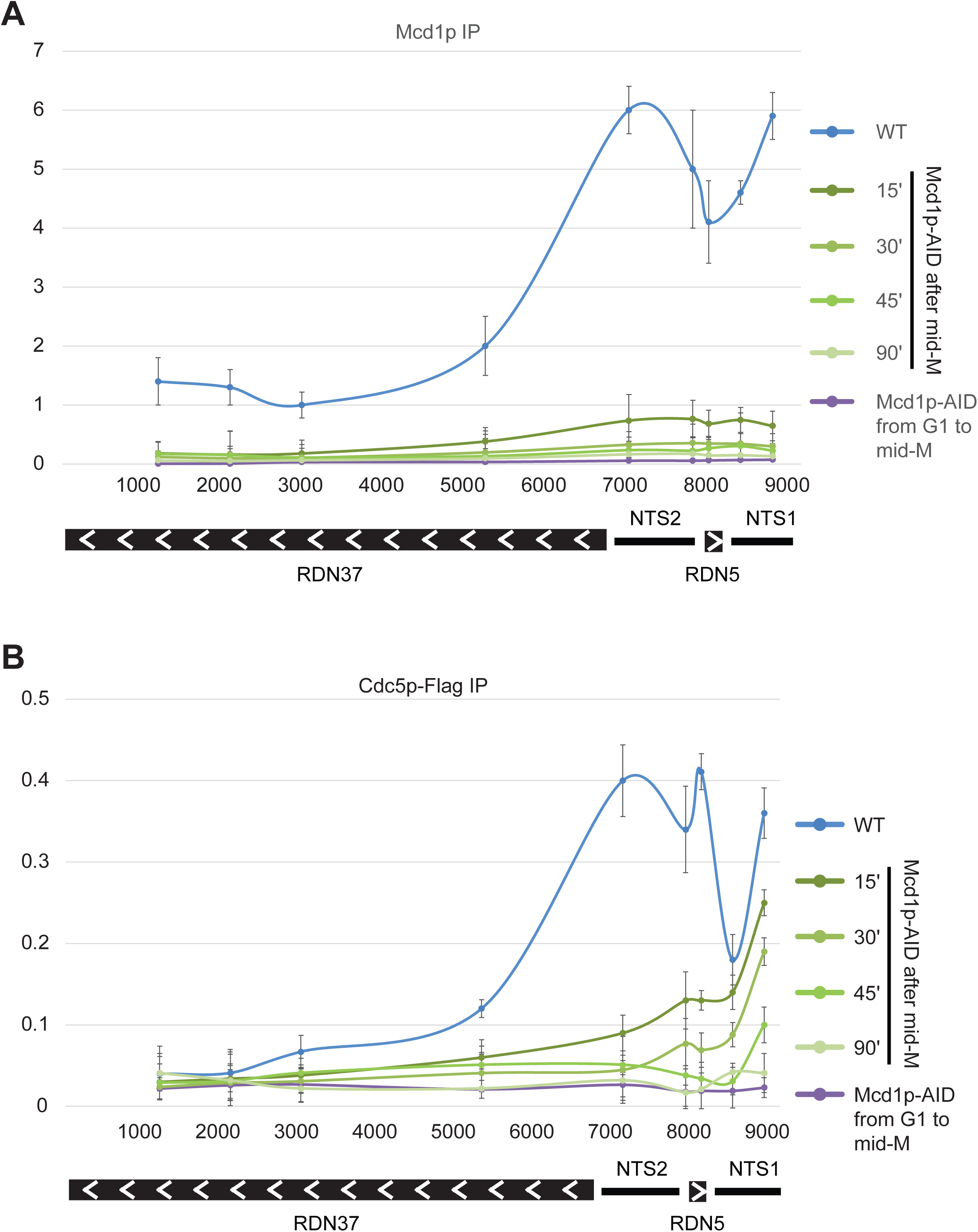
*Cdc5p and Mcd1p binding to RDN monitored by ChIP followed by quantitative PCR* **A.** Cohesin localization at the repetitive RDN locus. ChIP-seq followed by qPCR from a portion of chromosome XII are presented with lines. ChIP-qPCR signal for Mcd1p from WT cells arrested in mid-M is in blue, from cells depleted of Mcd1p-AID from G1 to mid-M and arrested in mid-M is in purple. Signal from cells arrested in mid-M and then depleted for Mcd1p-AID for the indicated time are in green. The region presented includes one copy of the rDNA repeat (9.1kb) made of the transcribed regions RND37 and RND5 (black box with white arrows indicating the direction of transcription) and the non-transcribed regions NTS1 and NTS2 (black line). Oligos sequences are indicated in q-PCR oligos table. **B.** Cdc5p localization at the repetitive RDN locus. ChIP-seq followed by qPCR from a portion of chromosome XII are presented with lines. ChIP-qPCR signal for Cdc5p from WT cells arrested in mid-M is in blue, from cells depleted of Mcd1p-AID from G1 to mid-M and arrested in mid-M is in purple. Signal from cells arrested in mid-M and then depleted for Mcd1p-AID for the indicated time are in green. The region presented includes one copy of the rDNA repeat (9.1kb) made of the transcribed regions RND37 and RND5 (black box with white arrows indicating the direction of transcription) and the non-transcribed regions NTS1 and NTS2 (black line). Oligos sequences are indicated in q-PCR oligos table.

**Figure 5—figure supplement 1:**
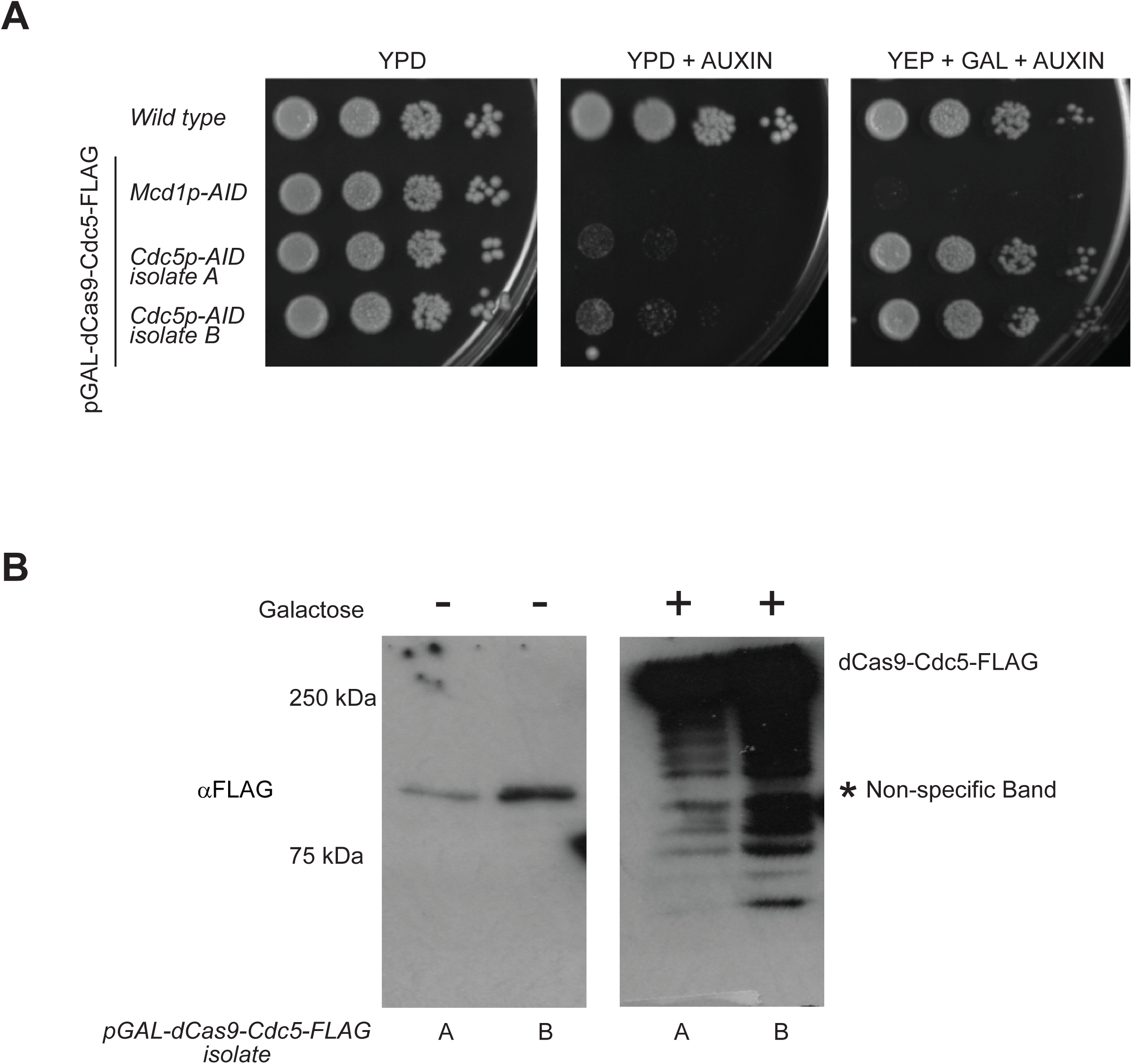
*Characterization of dCas9-Cdc5p strains.* **A.** dCas9-Cdc5p supports viability in the absence of endogenous Cdc5p. Asynchronous cells were serially diluted and plated on YPD (left), YPD with 750uM auxin (center), or YPD with 750uM auxin and 2% galactose (right). Plates were then incubated at 23C for three days and imaged. Wild Type, Mcd1p-AID with pGAL-dCas9-Cdc5p-FLAG, Cdc5p-AID with pGAL-dCas9-Cdc5p-FLAG isolate 1 and 2 were plated. **B.** Overexpression of dCas9-Cdc5p-FLAG by Western blotting.pGAL-dCas9-Cdc5p-FLAG isolate 1 and 2 strains were grown to mid-log in YPD. Cells were then split and grown either in the presence or absence of galactose for 1h. Cells were then pelleted and protein extracted by TCA as described in the materials and methods. Western blot analysis used FLAG antibody for dCas9-Cdc5p-FLAG.

**Figure 5—figure supplement 2:**
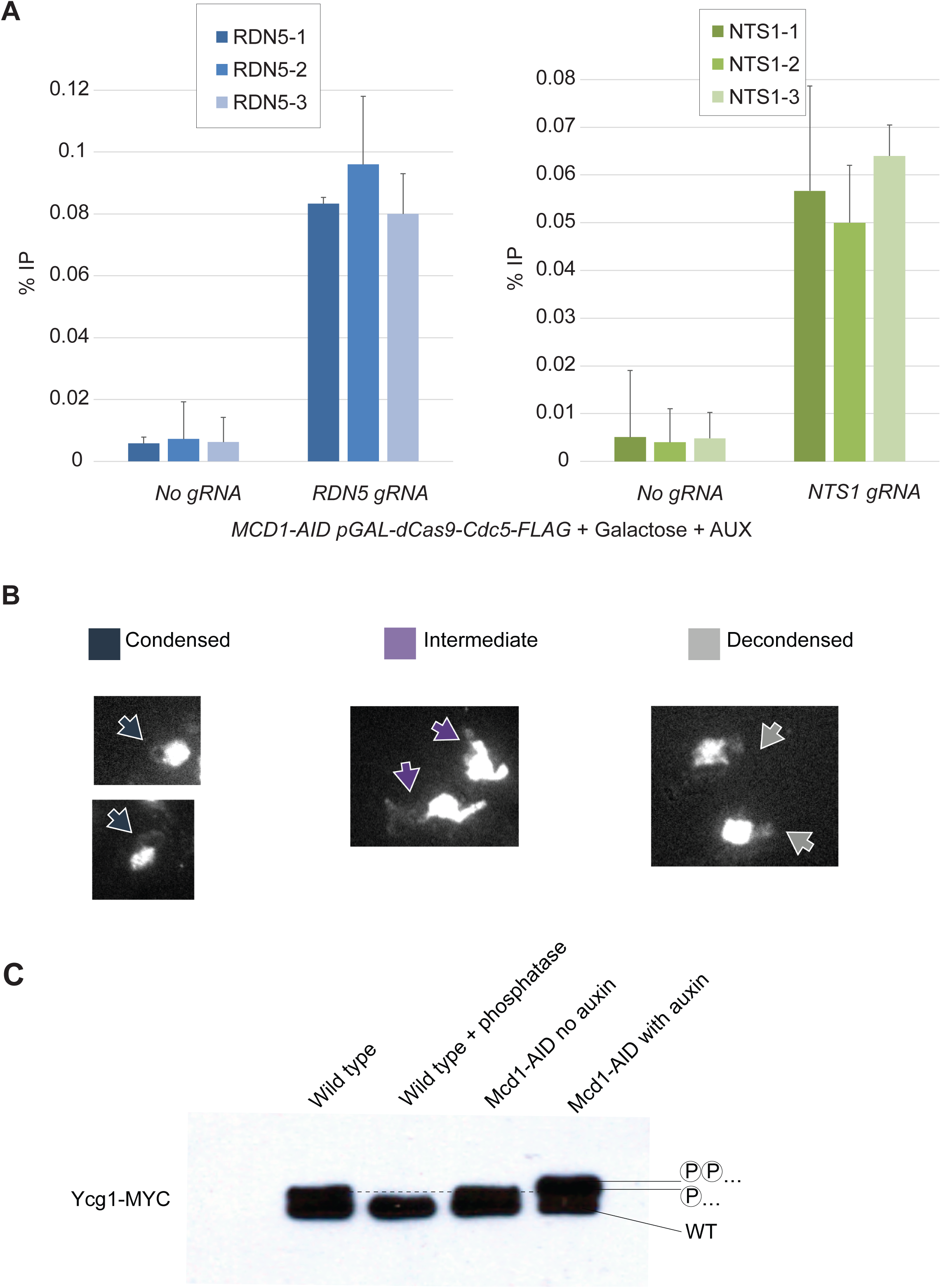
*Chromosomes spread pictures representing the different RDN morphologies observed upon dCas9-Cdc5p expression.* **A.** dCas9-Cdc5p is recruited to the RDN by gRNA. dCas9-Cdc5p-Flag binding to RDN5 ad NTS1 regions of the RDN are analyzed by ChIP followed by quantitative PCR in a strain depleted for Mcd1p-AID with auxin and induced for dCas9-Cdc5p-Flag with galactose. Signal coming from the RDN5 locus (3 different pairs of oligos were used) are presented in blu, from cells with no gRNA and cells expressing the gRNA for RDN5. Signal coming from the NTS1 locus (3 different pairs of oligos were used) are presented in green, from cells with no gRNA and cells expressing the gRNA for NTS1. Oligos sequences are indicated in q-PCR oligos table. **B.** dCas9-Cdc5p partially condenses RDN in the absence of endogenous Mcd1p. ‘Staged mid-M’ regimen: cultures were synchronized in G1 and released in media containing galactose and auxin to drive the expression of dCas9-Cdc5p concomitantly to the depletion of Mcd1p-AID. Media also contained nocodazole to arrest cells in mid-M and assess condensation. Mcd1p depleted with dCas9-Cdc5p and NTS1 gRNA expressing cells were subjected to staged mid-M’ regimen and processed to score RDN condensation. Representative images of chromosome spreads show a condensed loop of the RDN in the ‘condensed’ class, a partially condensed line in the ‘intermediate’ class, and a puffed RDN in the ‘decondensed’ class. **C.** Condensin (Ycg1p) is phosphorylated with a different pattern when cohesin is depleted. Phos-tag gel probed for condensin subunit Ycg1-Myc. Protein extract from a wild type strain (first lane), extract from wild type cells treated with phosphatase (second lane), from *MCD1-AID* strain not depleted (third lane), and from *MCD1-AID* strain depleted with Auxin (forth lane). The lower band represents the non-phosphorylated form of Ycg1p-Myc, above is the band with the pattern of phosphorylation observed in wild type cells (dotted line, one circled P), and above that is a third band with a different pattern of phosphorylation present just when Mcd1p-AID is depleted (two circled P).

## Materials and Methods

### Strain table

| Strain | Genotype | Reference |
| --- | --- | --- |
| DK5542 | MATa MCD1-AID-KANMX6 ADH1-OsTIR1- URA3::ura3-52 lys4::LacO(DK)-NAT trp1-1 GFPLacl-HIS3:his3-11,15 bar1 leu2-3,112 | Eng et al. 2014 |
| TE228 | MATa PDS5-3V5-AID2-KANMX6 lys4::LacO(DK)-NAT pHIS3-GFP-LacIHIS3::his3-11,15 trp1-1 ura3-52 | Eng et al. 2014 |
| RL06 | MATa CDC5-3V5-AID2-HPHMX ADH1-OsTIR1- URA3::ura3-52 lys4::LacO(DK)-NAT trp1-1 GFPLacl-HIS3:his3-11,15 bar1 leu2-3,112 | This study |
| RL401 | MATa BRN1-D375-3V5-AID2-HPHMX trp1Δ::pGPD1-TIR1-CaTRP1 lys4::LacO(DK)-NAT leu2-3,112 pHIS3-GFPLacl-HIS3:his3-11,15 ura3-52 bar1 | This study |
| VG3620-4C | MATa trp1Δ::pGPD1-TIR1-CaTRP1 lys4::LacO(DK)-NAT leu2-3,112 pHIS3-GFPLacl-HIS3:his3-11,15 ura3-52 bar1 | Çamdere et al. 2015 |
| RL404 | MATa CDC5-3V5-AID2-HPHMX YCG1-S406-6MYC-KANMX ADH1-OsTIR1- URA3::ura3-52 lys4::LacO(DK)-NAT trp1-1 GFPLacl-HIS3:his3-11,15 bar1 leu2-3,112 | This study |
| RL405 | MATa MCD1-AID-KANMX6 YCG1-S406-6MYC-HPHMX ADH1-OsTIR1- URA3::ura3-52 lys4::LacO(DK)-NAT trp1-1 GFPLacl-HIS3:his3-11,15 bar1 leu2-3,112 | This study |
| RL406 | MATa MCD1-AID-KANMX6 BRN1-D375-3V5-AID2-HPHMX ADH1-OsTIR1- URA3::ura3-52 lys4::LacO(DK)-NAT trp1-1 GFPLacl-HIS3:his3-11,15 bar1 leu2-3,112 | This study |
| RL407 | MATa MCD1-AID-KANMX6 CDC5-3V5-AID2-HPHMX ADH1-OsTIR1- URA3::ura3-52 lys4::LacO(DK)-NAT trp1-1 GFPLacl-HIS3:his3-11,15 bar1 leu2-3,112 | This study |
| RL410 | MATa MCD1-AID-KANMX6 CDC5-3FLAG-HPHMX ADH1-OsTIR1- URA3::ura3-52 lys4::LacO(DK)-NAT trp1-1 GFPLacl-HIS3:his3-11,15 bar1 leu2-3,112 | This study |
| RL411 | MATa BRN1-D375-3V5-AID2-HPHMX CDC5-3FLAG-KANMX ADH1-OsTIR1- URA3::ura3-52 lys4::LacO(DK)-NAT trp1-1 GFPLacl-HIS3:his3-11,15 bar1 leu2-3,112 | This study |
| RL413 | MATa CDC5-3V5-AID2-HPHMX<br>cdc5-K110A::LEU2 ADH1-OsTIR1- URA3::ura3-52 lys4::LacO(DK)-NAT trp1-1 GFPLacI-HIS3:his3-11,15 bar1 leu2-3,112 | This study |
| RL414 | MATa YCG1-S406-6MYC-KANMX trp1Δ::pGPD1-TIR1-CaTRP1<br>lys4::LacO(DK)-NAT leu2-3,112 pHIS3-GFPLacI-HIS3:his3-11,15 ura3-52 bar1 | This study |
| RL415 | MATa CDC5-3FLAG-HPHMX trp1Δ::pGPD1-TIR1-CaTRP1 lys4::LacO(DK)-NAT leu2-3,112 pHIS3-GFPLacI-HIS3:his3-11,15 ura3-52 bar1 | This study |
| RL416 | MATa MCD1-AID-KANMX6 TIR1-LEU2 pHIS3-GFPLacI-HIS3:his3-11,15 ura3-52 bar1 | This study |
| RL417 | MATa MCD1-AID-KANMX6 pGAL-dCas9-CDC5-3FLAG-TRP1:trp1Δ (RDN5 2u URA3) TIR1-LEU2 pHIS3-GFPLacI-HIS3:his3-11,15 ura3-52 bar1 | This study |
| RL418 | MATa MCD1-AID-KANMX6 pGAL-dCas9-CDC5-3FLAG-TRP1:trp1Δ (NTS1 2u URA3) TIR1-LEU2 pHIS3-GFPLacI-HIS3:his3-11,15 ura3-52 bar1 | This study |
| RL419 | MATa CDC5-AID-HPHMX TIR1-LEU2 pHIS3-GFPLacI-HIS3:his3-11,15 ura3-52 bar1 | This study |

### Reagents table

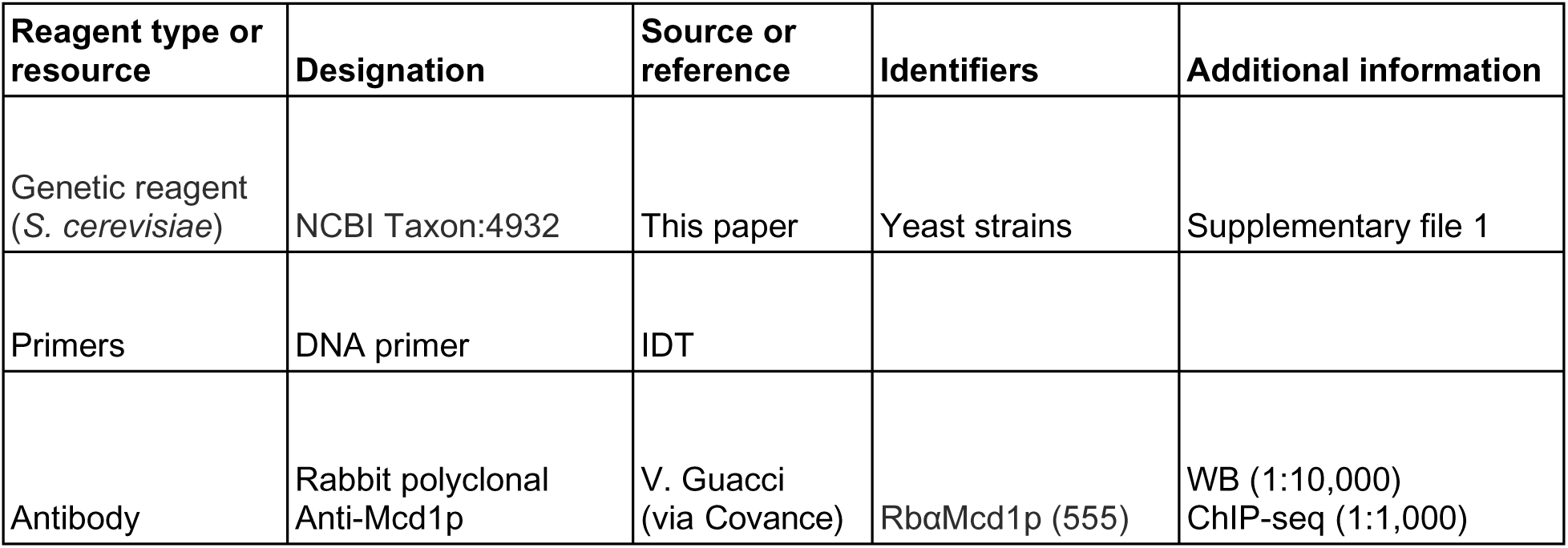
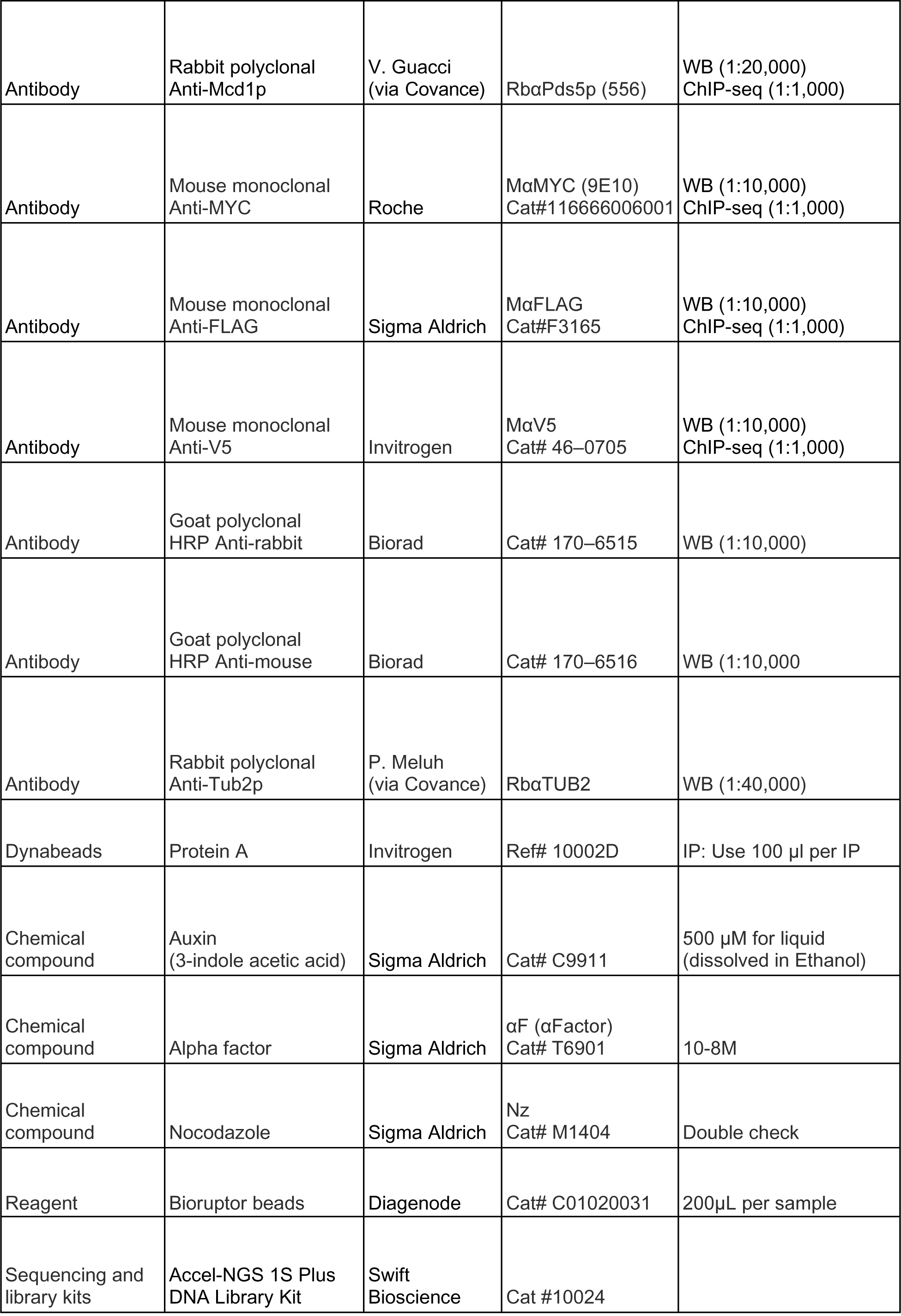

### q-PCR oligos table

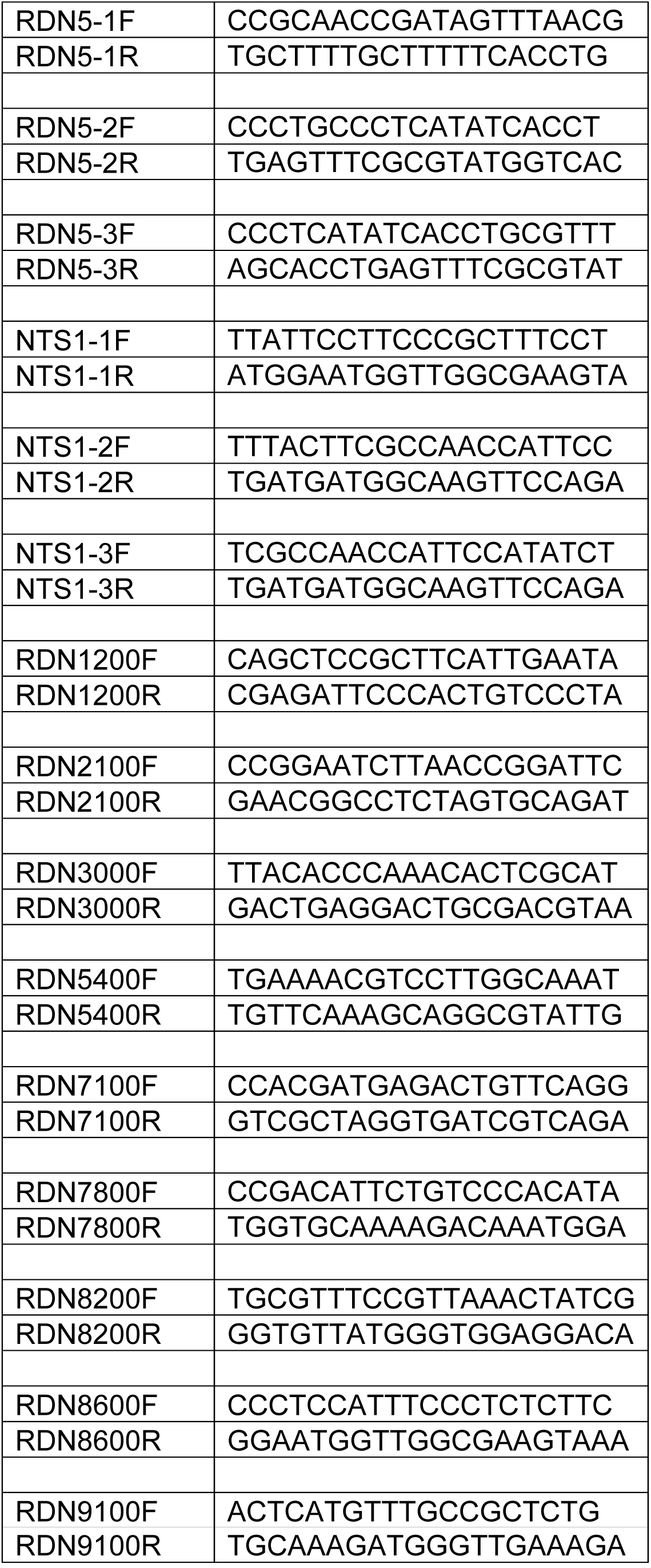

## Materials and Methods

### Yeast strains, growth and media

All strains used in this study are derived from the A364A background and their genotypes can be found in Table 1. YEP (yeast extract, peptone, dextrose) media and YPG (yeast extract, peptone and galactose) were prepared as previously described (Guacci 1997). Conditional AID degron strains were grown in YPD or YPG and auxin (3-indoleacetic acid; Sigma Aldrich Cat. I3750) at a final concentration of 500µM to deplete AID-tagged proteins. YPD auxin agar plates were prepared by cooling molten YPD 2% agar to 55°C and supplemented with auxin at a final concentration of 750µM.

### Sister chromatid cohesion

Sister chromatid cohesion was assessed as previously described (Robison and Koshland 2017). In brief, in our strains, a lacO array is integrated at the *LYS4* locus on chromosome IV. These arrays can be visualized by binding of GFP-lacI, which is integrated at the *HIS3* locus. To assess cohesion, cells were grown to early-log phase (OD600 0.1-0.2) at 23°C overnight and arrested in G1 using the pheromone alpha factor at a concentration of 10-8M (Sigma-Aldrich T6901-5MG). If indicated, auxin was added to a final concentration of 500µM to deplete AID-tagged proteins after a 2:30h arrest. Cells were released from G1 by washing with YPD containing auxin (if indicated) and 0.15mg/mL pronase E (Sigma Aldrich) three times, and once with YPD plus auxin. After the last wash Cells were resuspended in YPD medium containing auxin and 15 µg/mL nocodazole (Sigma Aldrich). Cultures were then once again incubated at 23°C for 3h. 1mL of these cells was then fixed in 70% ethanol and stored at 4C until ready for visualization. Cohesion was assessed by scoring the number of GFP-lacI foci in each cell on an Axioplan2 microscope (Zeiss, Thornwood, NY) using the 100X objective (numerical aperture 1.40) equipped with a Quantix charge-coupled camera (Photometrics). For time course experiments, 1mL of cells was collected every 15mn after release from alpha factor.

### Assessing condensation at the RDN locus

Cells were grown as described for the sister chromatid cohesion assay. After 3h in nocodazole, cells were fixed, spheroplasted and prepared for FISH as previously described (Guacci et al 1994). DNA masses were then visualized with DAPI (4’,6-diamidino-2-phenylindole) and imaged as described above. RDN morphology was scored as condensed “looped” or decondensed “puffy” as previously described (Guacci et al. 1994).

### Maintenance Assay

For maintenance assays, cells were grown to early log phase at 23C and arrested in mid-M using 15µg/mL nocodazole. For AID-tagged proteins, auxin was added after synchronous mid-M arrest, 2h after nocodazole addition. Cells were collected after 1h of auxin addition for condensation assay.

### Add-later time course

Cells were grown as described in the sister chromatid cohesion assay to get synchronous population of cells arrested in mid-M depleted for the AID-tagged protein(s). To restore the presence of the AID-tagged proteins, these cells were washed 3 times in YPD medium containing nocodazole and grown in media containing nocodazole for 2h. Samples were collected after 2h to assess condensation state and protein levels by TCA extraction followed by Western.

### TCA protein extraction and Western blotting

To assess protein level, cells were collected, pelleted, washed with 1X PBS and frozen at −80. Frozen pellets were resuspended in 200uL 20% TCA and broken for 40s using a FastPrep-24 5G instrument (6.0m/sec each) (MP Bio, Santa Ana, CA). Lysates were diluted with 1mL 5%TCA and spun for 10mn at 14,000rpm at 4C. Pellets were resuspended in 2X Laemmli buffer and boiled for 7mn and spun. Cleared lysates were used for Western blotting on 6% SDS-PAGE gels. Phos-tag gels were prepared, run and transferred following manufacturer protocol (alphalabs).

### Chromatin immunoprecipitation (ChIP) and sequencing

Cells were grown and arrested at 30oC as described for assessing condensation. 60OD of mid-M cells were collected, fixed for one hour with 1% formaldehyde, quenched with 5mM glycine and processed for ChIP as previously described (Robison and Koshland 2017, with the following modifications). Frozen cell pellets were disrupted 3 times using a FastPrep-24 5G instrument (40 seconds, 6.0m/sec each) (MP Bio, Santa Ana, CA). Chromatin was sheared 10 x 30s ON/30s OFF with a Bioruptor Pico (Diagenode, Denville, NJ). For ChIP-qPCR, 1/20 of sheared and cleared lysate was reserved as input. Chromatin extracts were incubated with 4uL of the following antibodies monoclonal mouse anti-MYC (Roche), monoclonal mouse anti-FLAG (Sigma Aldrich), polyclonal rabbit anti-Pds5p (Covance Biosciences, Princeton, NJ), or polyclonal rabbit anti-Mcd1p (Covance Biosciences, Princeton, NJ) overnight at 4C. Antibody-bound lysates were incubated with 120uL Protein A dynabeads (Invitrogen, Cat#: 10001D) for 1h. Following ChIP, the library was prepared using the Accel-NGS 1S Plus DNA Library Kit (Swift Bioscience) following the manufacturer protocol. Libraries were sequenced using Illumina Hiseq 4000. The sequencing files were aligned to the SacCer3 yeast genome using Bowtie2 tool and bigwig files were normalized for number of reads.

### Accession numbers

The data generated are available at Gene Expression Omnibus (GEO) under accession numbers GSE147290

## dCas9-Cdc5p experiments

Cells were grown in 5mL of -URA media until ready to be used to maintain plasmid. Cells were grown to early-log phase (OD600 0.2) at 23°C overnight and arrested in G1 using the pheromone alpha factor at a concentration of 10-8M (Sigma-Aldrich T6901-5MG) in YEP+ 2% lactic acid medium. After 2:30h of growth in alpha factor, auxin was added to a final concentration of 500µM to deplete AID-tagged proteins and galactose was added to a final concentration of 2% to induce expression of pGal-dCas9-Cdc5-3Flag construct. After an hour, cells were released from G1 by washing with YEP+2% Galactose medium containing auxin and 0.15mg/mL pronase E (Sigma Aldrich) three times, and once with YEP+2% Galactose plus auxin. After the last wash, cells were resuspended in YEP+2% Galactose medium containing auxin and 15 µg/mL nocodazole (Sigma Aldrich). Cultures were then once again incubated at 23°C for 3h. 1mL of these cells was then processed as described in the “assessing condensation at the RDN locus” methods.

## Acknowledgements

We thanks Thomas Eng and Vincent Guacci for building some of the strains used. We thank Elcin Unal for sharing the dCas9 plasmid and Fred Winston for critical reading of the manuscript. We thank the Koshland lab and the Unal lab for imputs and discussions. This work was funded by National Science Foundation GRFP grant to RL and National Institutes of Health Grant (1R35 GM-118189-01 to DK).

